# The Influenza B Virus Victoria and Yamagata Lineages Display Distinct Cell Tropism and Infection Induced Host Gene Expression in Human Nasal Epithelial Cell Cultures

**DOI:** 10.1101/2023.08.04.551980

**Authors:** Jo L. Wilson, Elgin Akin, Ruifeng Zhou, Anne Jedlicka, Amanda Dziedzic, Hsuan Liu, Katherine Z. Fenstermacher, Richard Rothman, Andrew Pekosz

## Abstract

Understanding Influenza B virus infections is of critical importance in our efforts to control severe influenza and influenza-related disease. Until 2020, two genetic lineages of influenza B virus – Yamagata and Victoria – circulated in the population. These lineages are antigenically distinct but differences in virus replication or the induction of host cell responses after infection have not been carefully studied. Recent IBV clinical isolates of both lineages were obtained from influenza surveillance efforts of the Johns Hopkins Center of Excellence in Influenza Research and Response and characterized *in vitro*. B/Victoria and B/Yamagata clinical isolates were recognized less efficiently by serum from influenza-vaccinated individuals in comparison to the vaccine strains. B/Victoria lineages formed smaller plaques on MDCK cells compared to B/Yamagata, but infectious virus production in primary human nasal epithelial cell (hNEC) cultures showed no differences. While ciliated epithelial cells were the dominant cell type infected by both lineages, B/Victoria lineages had a slight preference for MUC5AC-positive cells, while B/Yamagata lineages infected more basal cells. Finally, while both lineages induced a strong interferon response 48 hours after infection of hNEC cultures, the B/Victoria lineages showed a much stronger induction of interferon related signaling pathways compared to B/Yamagata. This demonstrates that the two influenza B virus lineages differ not only in their antigenic structure but in their ability to induce host innate immune responses.

## Introduction

Circulating influenza viruses are members of the family, Orthomyxoviridae. There are four types of human influenza viruses, with Influenza A Virus (IAV) and Influenza B Virus (IBV), resulting in the majority of clinically important infections [1]. The CDC estimates that between 2010 and 2020, seasonal influenza viruses resulted in 9-41 million illnesses, 140,000-710,000 hospitalizations and 12,000-52,000 deaths as of Feb 3, 2023 [2]. Research on seasonal influenza is biased towards IAV due to IAV accounting for the majority of annual infections. IAV, but not IBV, also poses a pandemic threat due to its ability to infect a wide variety of animal species. IBV is not known to have an animal reservoir and circulates exclusively in humans [3]. Prior to the SARS-CoV-2 pandemic, IBV represented 20% of confirmed influenza cases and is consistently underestimated as to its impact on healthcare burden [4], [5]. IAV and IBV infection leads to similar acute clinical syndromes with a wide range of severity [6], [7]. Although clinical severity in the general population appears similar across studies, there have been reports of higher IBV-attributable mortality in children and individuals infected with HIV [5], [8].

Circulating IBV is divided into two antigenically distinct lineages defined by the Hemagglutinin (HA) genomic segment, B/Victoria/2/1987 like (B/Victoria) and B/Yamagata/16/1988-like (B/Yamagata). In 2012, it was first recommended by the FDA to include both lineages of IBV in a quadrivalent vaccine to replace the trivalent vaccines containing only one IBV lineage [9]. The recommendation was made due to shifting patterns of dominance between IBV lineages resulting in frequent lineage level mismatch as well as limited cross-protection [5], [10]–[13]

Numerous reports in the literature show that these two virus lineages behave differently in the population suggesting unique features of each that deserve further research. The primary differences noted are patterns of evolutionary escape, variation in predominance based on temperate or tropical climate and age predilection. After 2010, B/Yamagata viruses diversified into multiple, coexisting clades followed by the dominance of a single clade, clade 3, in 2016. B/Yamagata drift has been defined not only by single nucleotide changes in HA but more so driven by single nucleotide changes in Neuraminidase (NA). B/Victoria viruses have had high antigenic diversity since 2010 with frequent insertion and deletion events in the HA receptor binding site leading to immune evasion, a strategy observed in B/Victoria evolution since its first isolation [14], [15]. There is also a clear difference in age predisposition between these two virus lineages [4], [5], [16]. B/Yamagata viruses have a higher average age of infection commonly infecting older adults, whereas B/Victoria viruses show high pediatric rates of infections. One of the highest rates of pediatric mortality attributable to influenza occurred during the 2019-20 Northern Hemisphere influenza season where 61% of pediatric deaths were attributable to B/Victoria lineage despite making up only 41% of total infections (accessed February 3, 2023) [17]. These patterns are well described for IBV, however, the mechanisms that drive these differences remain unknown highlighting the importance of research to define differences in these virus lineages.

Literature detailing the respiratory epithelium response to IBV infection is limited. Bui et al published a detailed study of cell tropism and replication kinetics of B/Yamagata and B/Victoria lineage viruses. They showed IBV replication in all human airway organoid cell types (ciliated, mucus-containing, secretory, and basal cells) using immunohistochemistry at 24- and 48-hours post-infection (hpi). Using real-time PCR, they evaluate CCXL10, IFN-β, and CCL5 between isolates of B/Yamagata and B/Victoria showing minor strain level differences with no significant lineage level variation. Additionally, clinical evaluations of serum cytokines have shown that IL-17A, IFN-g, and IP-10 are dominant responses after IBV clinical infections [18]. We sought to define IBV–epithelial cell interactions including viral kinetics, and quantitative cell tropism over the full course of infection using flow cytometry, as well as protein immunoassays and transcriptomics using human primary, differentiated nasal epithelial cell cultures.

## Materials and Methods

### IBV Clinical Isolate Collection from Johns Hopkins Hospital, Baltimore, MD

The human subjects’ protocol was approved by the Johns Hopkins School of Medicine Institutional Review Board (IRB90001667) and the National Institutes of Health Division of Microbiology and Infectious Diseases (protocol 15-0103). Patients were enrolled at the Johns Hopkins Medical Institute (JHMI) Department of Emergency Medicine or on inpatient floors. Symptomatic patients in the emergency department were screened and tested for influenza from triage by clinical providers using a validated clinical decision guideline tool. After written consent was obtained, a nasopharyngeal swab was obtained and stored at -80°C.

### Cell Lines and Cell Culture Maintenance

Madin-Darby Canine Kidney Cells (MDCK) and MDCK cells overexpressing 2,6 sialyltransferase (MDCK-SIAT-1) were maintained in complete medium (CM) consisting of Dulbecco’s Modified Eagle Medium (DMEM) supplemented with 10% fetal bovine serum, 100U/mL of penicillin and 100ug/mL of streptomycin mixture (Life Technologies) and 2mM Glutamax (Gibco). Primary Human Nasal Epithelial Cells (Promocell, Heidelberg, Germany) were plated and cultured in Ex-Plus Medium (Stem Cell Technologies, Pneumacult Ex-Plus Media Kit) without antibiotics. The apical surface of the wells was coated with 0.03 mg/mL Collagen I, Rat Tail (Gibco). One tube of cryopreserved cells (∼500,000) is then directly plated on a 24-well transwell plate, with cells divided equally between 24-6.5 mm 0.4uM wells with Ex-Plus media on the apical and basolateral surface. This media promotes proliferation and inhibits terminal differentiation. The media is changed 24 hours after cell plating and every 48 hours following to maintain cell viability. After 7-10 days, confluence is assessed using visual monitoring by microscopy and objectively measured by Transepithelial Electrical Resistance (TEER). When cell monolayers reached a TEER greater or equal to 400Ω-cm2, both apical and basolateral media were removed and ALI Differentiation media (Stem Cell Technologies, Pneumacult ALI Basal Medium supplemented with 1X ALI Maintenance Supplement (StemCell Technologies), 0.48 ug/mL Hydrocortisone solution (StemCell Technologies), and 4 ug/mL Heparin sodium salt in PBS (StemCell Technologies) was replaced on the basolateral side only. This change allows full differentiation of human nasal cultures. Media is changed every 48 hours to maintain cell viability. The apical surface of cells is intermittently washed with PBS to remove excess mucus. Full differentiation takes approximately 4 weeks and cells are considered fully differentiated when there is presence of mobile cilia on the cell surface visible with light microscopy. Cells are used for experiments once considered fully differentiated and remain viable for 4-6 weeks following differentiation. All cells were maintained at 37°C in a humidified incubator supplemented with 5% CO2. Influenza B virus infections of hNEC cultures were carried out at 33°C to model nasopharynx temperature. Plates are placed at 33°C 24 hours prior to infection to acclimate cells to infection temperature.

### IBV Lineage Determination-RT PCR

Nasal swabs from the 2016-2017, 2017-2018, and 2019-2020 seasons that tested positive for IBV rapid PCR in the JH CEIRS database were used. We chose a two-step lineage determination process for IBV-positive PCR nasopharyngeal swabs. Lineage is also then confirmed with HA gene segment sequencing. We used WHO recommended line-age-specific primers and using RT-PCR, generated oligonucleotides of specific sizes based on lineage [19]. For viruses collected post-2017, we used updated primers to account for observed hemagglutinin deletions at positions 162-164 [20]. Viral RNA was isolated using Qiagen QIAamp Viral RNA Mini Kit per the manufacturer’s protocol. 140µl of nasopharyngeal swab sample was used for each extraction. The concentration of extracted vRNA was measured by NanoDrop and 2µl of vRNA were input into RT-PCR reactions. One-step RT-PCR master mix was prepared with SuperScript™ III One-Step RT-PCR System with Platinum™ Taq DNA Polymerase per the manufacturer’s instruction. All 4 primers were added to the mix at a final concentration of 10uM.

**Table 1.**
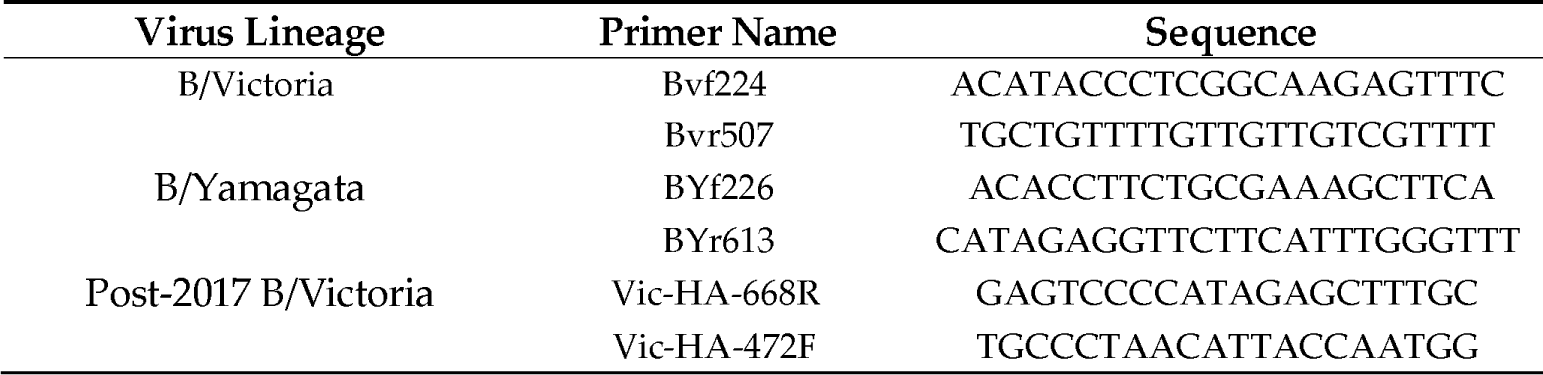
Primers for Lineage Determination.

### Virus isolation

Nasopharyngeal swabs or nasal wash originating from B/Victoria or B/Yamagata positive individuals were used for virus isolation on primary cells. The hNEC cultures were washed three times with 150ml of phosphate buffered saline (PBS, Gibco) containing 0.9mM Ca2+ and 0.5mM2+ (PBS+), with incubations of 10 minutes at 37°C for each wash. 150ml of the nasal swab sample was then placed on the apical side of the hNEC cultures and samples incubated at 37°C for 2 hours. Following incubation, the sample was aspirated, and the cells were washed twice with PBS+. Cultures were then incubated at 37°C. On days 3-, 5-, and 7-days post-infection, 150ml of infectious media containing Dulbecco modified Eagle medium (Sigma), 10% penicillin/streptomycin (Gibco), 10% L-glutamine (Gibco), and 0.5% BSA (Sigma) was placed on the apical surface of the cells and incubated for 10 minutes at 37°C. The apical wash was harvested and stored at –65°C, followed by assessment of infectious virus by TCID50 assay.

### TCID50

MDCK-SIAT-1 cells were seeded in a 96-well plate 2 days before assay and grown to 100% confluence. Cells were washed twice with PBS+ then 180µL of IM was added to each well. Ten-fold serial dilutions of virus from 10-1 to 10-7 were created and then 20µL of the virus dilution was added to the MDCK-SIAT-1 cells. Cells were incubated for 6 days at 33°C then fixed with 2% formaldehyde. After fixing, cells were stained with naphthol blue-black, washed and virus titer was calculated using the Reed and Muench method [21].

For clinical isolates, the earliest sample that showed presence of infectious virus was used to infect a T75 Flask at an MOI of 0.01. Working stocks for each clinical isolate were generated by infecting a T75 flask of MDCK-SIAT-1 cells at an MOI of 0.001 for one hour at room temperature while rocking. The inoculum was removed, and cells were placed in a 33°C incubator and monitored daily for CPE. Working stock was harvested between 3 and 5 days or when CPE was seen in 75-80% of the cells. Harvested media was centrifuged at 400 xg for 10 minutes at 8°C to remove cell debris, and the resulting supernatant was aliquoted into 500µl and stored at -80°C infectious virus quantity of working stocks was determined using TCID-50 assay. Seed and working stocks of vaccine strains of IBV were grown directly in MDCK-SIAT-1 cells as described above. IBV vaccine strains were kindly provided by Johns Steel, Centers for Disease Control (CDC).

**Table 2.** Viruses Used in Comparison Studies. Virus seed and working stocks.

| Virus Lineage/Clade | Source | Virus Name | GISAID Accession |
| --- | --- | --- | --- |
| Victoria/V1A | Clinical isolate | B/Baltimore/R0122/2016 | EPI_ISL_17412143□ |
| Victoria/V1A | Clinical isolate | B/Baltimore/R0001/2016 | EPI_ISL_17412142 |
| Yamagata/Clade 3 | Clinical isolate | B/Baltimore/R0250/2018 | EPI_ISL_17412144 |
| Yamagata/Clade 3 | Clinical isolate | B/Baltimore/R0337/2018 | EPI_ISL_17412145 |
| Yamagata/Clade 3 | Clinical isolate | B/Baltimore/R0300/2018 | EPI_ISL_17742639□ |
| Victoria/V1A.1 | Vaccine (CDC) | B/Colorado/06/2017 | EPI_ISL_257735 |
| Victoria/V1A.3 | Clinical isolate | B/Baltimore/R0696/2019 | EPI_ISL_17353886 |
| Yamagata/Clade 3 | Vaccine (CDC) | B/Phuket/3073/2013 | EPI_ISL_161843□ |

### HA Sequencing and Lineage Assignment

Vaccine virus HA segments were sequenced to confirm that no sequence alteration occurred with the creation of working stocks. 100% sequence identity was maintained using the GISIAD sequence database for sequence comparison. Viral RNA from each sample was isolated using Qiagen vRNA isolation Kit. Superscript III RT-PCR system (ThermoFisher) was used to isolate cDNA for sequence analysis. Primers were designed at the 5’ and 3’ noncoding regions HA segment with the goal of sequencing the entire segment.

**Table 3.**
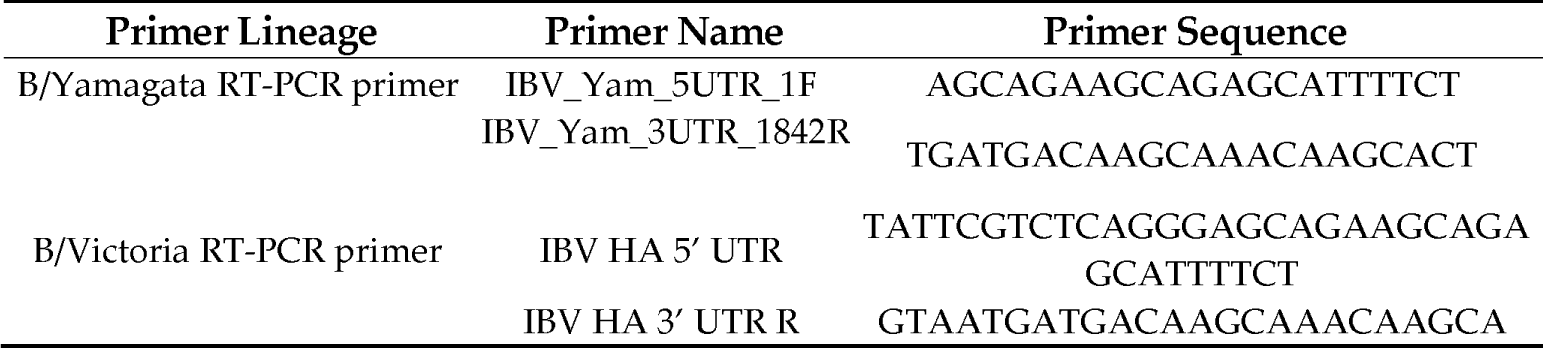
Primer Design. ^1^Tables may have a footer.

Additional sequencing primers were designed within the reading frame to ensure full coverage of the entire segment. The cDNA was sent to the Synthesis & Sequencing Facility of the Johns Hopkins University for Sanger sequencing.

### Influenza B Genome Sequencing

Viral RNA was extracted using the QIAamp viral RNA mini extraction kit. For library preparation, the Illumina RNA Prep with Enrichment (L) Tagmentation kit with Respiratory Virus Oligo Panel v2 (20044311) was used following the single-plex enrichment protocol, and samples were sequenced using a MiSeq Illumina sequencer (v3, 2×300 bp or 2× 75bp). Consensus sequences were generated using the DRAGEN RNA Pathogen Detection pipeline using custom .bed files and FASTA files for IBV.

### HA and NS Phylogenetics Analysis

Representative IBV HA and NS sequences between the years 2009 and 2022 were accessed from GISAID filtered to include only complete sequences [22]. Associated GISAD metadata for HA and NS sequences analyzed in this study are compiled in Supplementary Table 4. Sequences were aligned using MAFFT v7.520. For the HA tree, amino acid changes were standardized using HA numbering using the FluDB conversion tool. Alignments were used to construct a maximum likelihood time-resolved tree in treetime using a relaxed molecular clock model. Clade assignments for B/Victoria and B/Yamagata were assigned using NextClade v2.11.0 with the Influenza B lineage workflows. Constructed trees were annotated by lineage and clade in R v4.1.1 using ggtree v3.16 [23], [24].

### Neutralizing Antibody Assays

Serum samples used for this analysis were from Johns Hopkins Medical Institute healthcare workers recruited from the Johns Hopkins Centers for Influenza Research and Surveillance (JHCEIRS) during the annual employee influenza vaccination campaign in the Fall of 2019. Pre-vaccine and approximately 28-day post-serum samples were used for analysis. Subjects provided written informed consent prior to participation. The JHU School of Medicine Institutional Review Board approved this study, IRB00288258. Serum samples were first treated (1:3 ratio serum to enzyme) with Receptor Destroying Enzyme (Denka-Seiken) and incubated overnight at 37°C followed by inactivation at 57 °C for 35 minutes. Serum was diluted 2-fold in IM (Dulbecco modified Eagle medium (Sigma), with 10% penicillin/streptomycin (Gibco), 10% L-glutamine (Gibco), 0.5% BSA (Sigma), and 5µg/mL of N-acetyl trypsin (Sigma) at 37°C and 5% CO2) and 100 TCID50 was added for a one-hour incubation at room temperature. Serum Sample/Virus was used to infect a confluent layer of MDCK-SIAT-1 cells. The inoculums were removed after 24 hours, and fresh media (same recipe as above) was added for 96 hours. Plates were fixed and stained as described previously. The Neutralizing Antibody titer was calculated using the highest serum dilution that led to greater than 50% CPE.

### Plaque Assays

MDCK cells were grown in complete medium to 100% confluency in 6-well plates. Complete medium was removed, cells were washed twice with PBS containing 100µg/ml calcium and 100µg/ml magnesium (PBS+) and 250µL of inoculum was added. Virus dilution was done by serially diluting the virus stock 10-fold each time until 10-6, Cells were incubated at 33°C for 1 hour with rocking every 10 minutes. After 1 hour, the virus inoculum was removed and phenol-red free MEM supplemented with 3% BSA (Sigma), 100U/mL of penicillin and 100ug/mL of streptomycin mixture (Life Technologies), 2mM Glutamax (Gibco), and 5µg/ml N-acetyl trypsin (Sigma), 5mM HEPES buffer and 1% agarose was added. Cells were incubated at 33°C for 3-5 days and then fixed with 4% formaldehyde. After removing the agarose, cells were stained with napthol-blue black. Plaque size was analyzed in Image J.

### Low-MOI Infections

Low-MOI growth curves were performed at an MOI of 0.01 in hNEC cultures. The hNECs were acclimated to 33°C for 48 hours before infection. The apical surface was washed three times with PBS and the basolateral media was changed at time of infection. hNEC cultures were inoculated at an MOI of 0.01. hNEC cultures were then placed in a 33°C incubator for 2 hours. After incubation, the apical surface of the hNEC culture was washed three times with PBS+. At the indicated times, 100µl of IM without N-acetyl trypsin was added to the apical surface of the hNECs for 10 minutes at 33°C, the IM was harvested and stored at -80°C. Basolateral media was changed every 48 hours post infection for the duration of the experiment. Infectious virus titers in the apical supernatants were measured with TCID50 assay.

### Flow CytometryD

For the 72-hour time point data experiments: hNEC cultures were infected with IBV clinical isolates at an MOI of 0.01 for 72 hours. At 72 hours, cells were harvested from the apical membrane into a single cell suspension with a 30-minute incubation in 1X TrypLE. After cells were trypsinized, they were resuspended in a trypsin stop solution. The cells were then washed three times in 1X PBS and resuspended in 1 mL PBS (centrifuge at 400 xg between wash steps). Appropriate control and sample tubes were then stained with AQUA viability dye 1 uL/1×106 cells for 30 minutes at RT. Cells were then washed and resuspended in BD Fixation/Permeabilization solution and incubated for at least 30 minutes at 4°C. Cells were washed with BD Perm/Wash Buffer x2 and centrifuged at 400 xg at 4°C for 5 minutes. Cells were then resuspended in BD Perm/Wash Buffer with 7% NGS and incubated for 1 hour at 4°C. Cells were washed with BD Perm/Wash Buffer x2 and centrifuged at 400 xg at 4°C for 5 minutes. Appropriate sample tubes were incubated with primary antibodies for one hour at RT. Antibodies are diluted into BD Perm/Wash buffer at appropriate concentrations. Final staining volume is 200 ml. Cells were washed with BD Perm/Wash Buffer x2 at 400xg and centrifuged at 4°C for 5 minutes. Appropriate sample tubes were incubated with secondary antibodies for 30 minutes at RT. Cells were washed with BD Perm/Wash Buffer x2 at 400 xg and centrifuged at 4°C for 5 minutes. Appropriate sample tubes were incubated with conjugated antibodies for 30 minutes at RT. Cells were washed with BD Perm/Wash Buffer x2 and centrifuged at 400xg at 4°C for 5 minutes. Cells were resuspended in FACS Buffer and filtered through a 35 uM strainer cap into FACS tubes just prior to the run. Cell suspensions were run on a BD LSRII Flow Cytometer using DIVA software. Single stained cells were used as controls and fluorescence minus one controls were used to assist in gating. Other controls included secondary antibody alone controls and uninfected IBV HA stained controls to ensure there was no non-specific staining. Data analysis was completed on FlowJo v10. Gating strategy employed was as follows: exclusion of debris, single cells, and Aqua – cells (LIVE). For the time course experiment: Low MOI infection was completed at an MOI of 0.01. At each respective time point (72 hours post infection, 96 hours post infection and 120 hours post infection), cells were trypsinized, stained for live/dead discrimination and fixed in 4% Formaldehyde as detailed above. The samples were held at 4°C after fixation until collection of all samples. Samples were then blocked and stained all together exactly as detailed above and run on the LSRII.

**Table 4.**
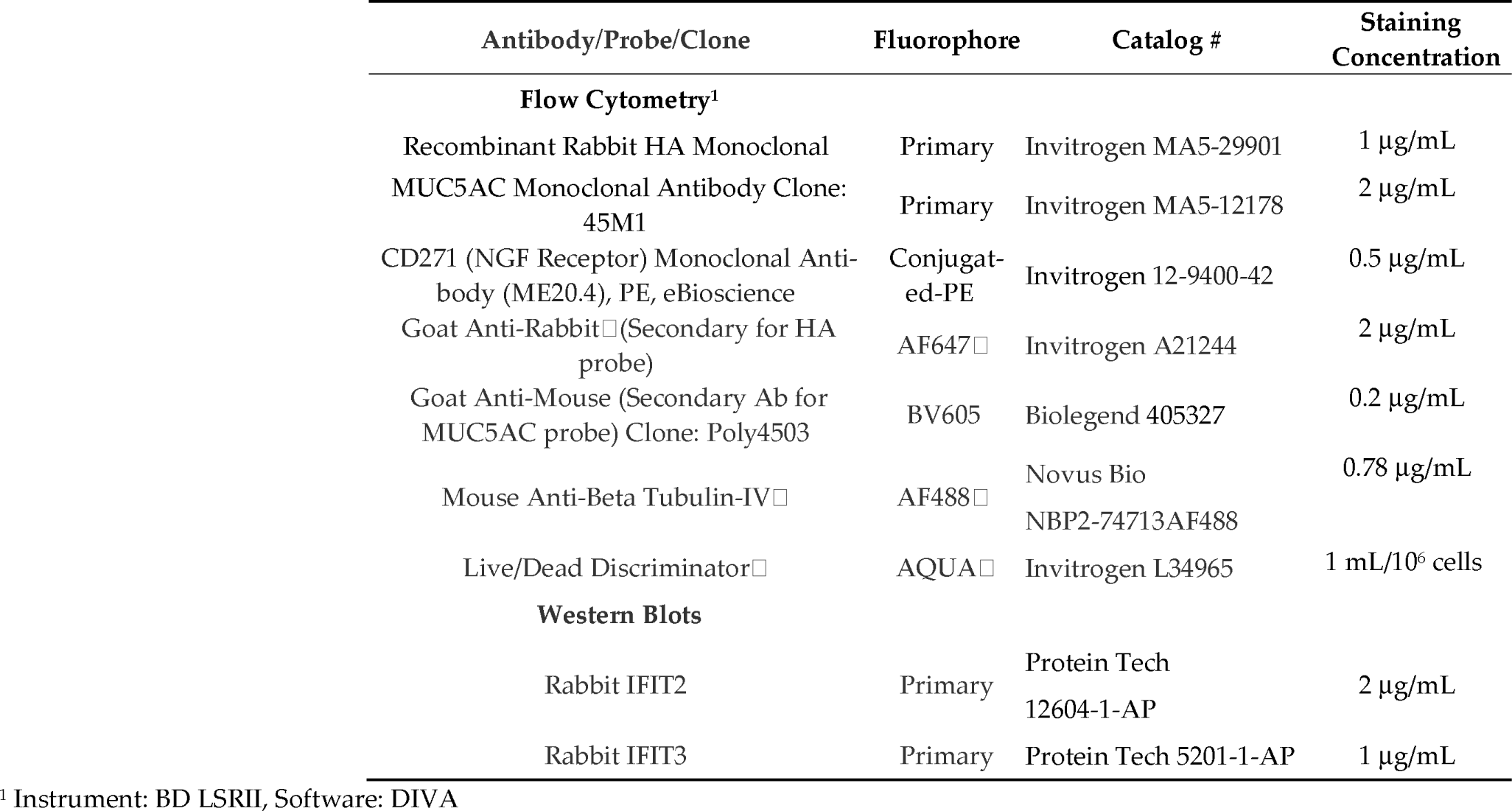
Antibody List.

**Table 5.**
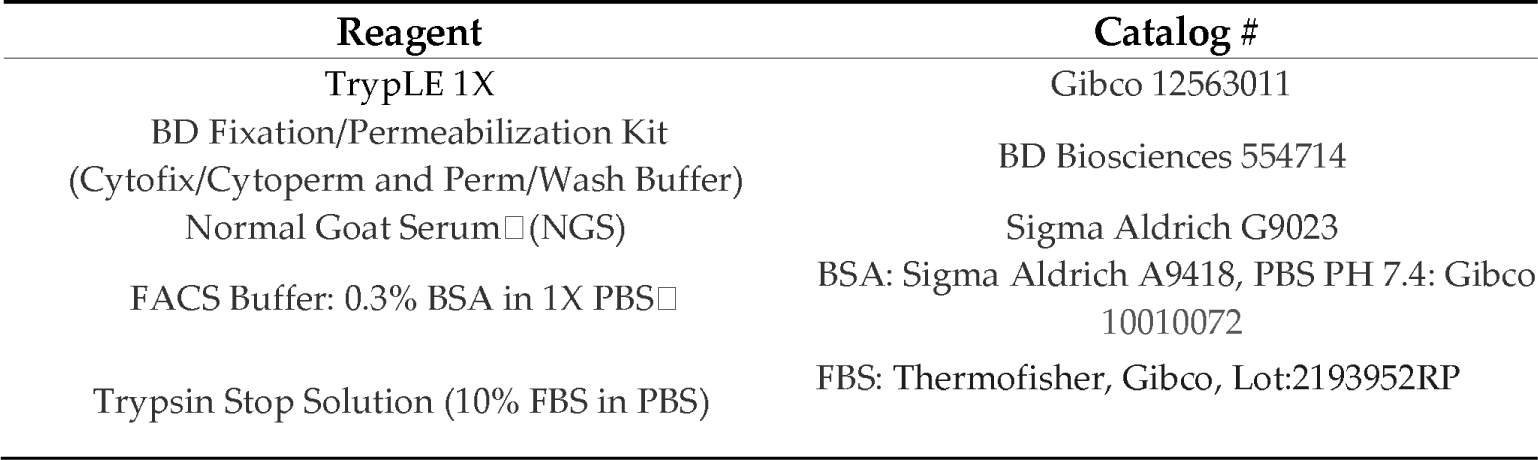
Flow Cytometry Reagent List.

| Reagent | Catalog # |
| --- | --- |
| TrypLE 1X | Gibco 12563011 |
| BD Fixation/Permeabilization Kit<br>(Cytofix/Cytoperm and Perm/Wash Buffer) | BD Biosciences 554714 |
| Normal Goat Serum (NGS) | Sigma Aldrich G9023 |
| FACS Buffer: 0.3% BSA in 1X PBS | BSA: Sigma Aldrich A9418, PBS PH 7.4: Gibco 10010072 |
| Trypsin Stop Solution (10% FBS in PBS) | FBS: Thermofisher, Gibco, Lot:2193952RP |

**Table 6.**
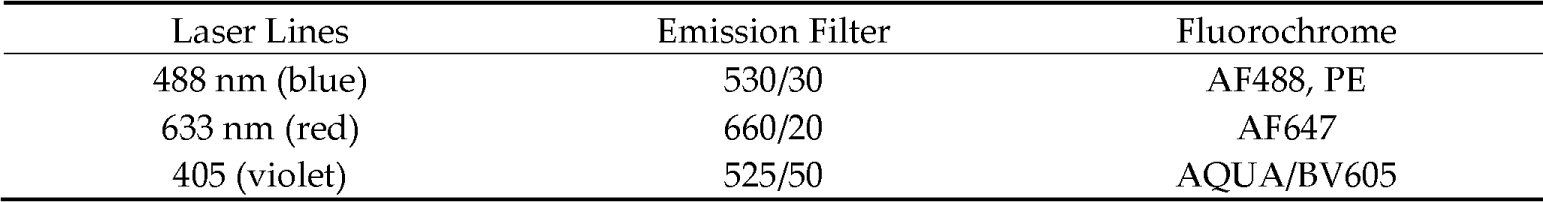
BD LSRII Set Instrument Set Up.

To evaluate cytokine and chemokine responses to infection, a custom Luminex panel was built including the following analytes: BAFF, E-Cadherin, Eotaxin-3, Eotaxin, G-CSF, IFN-a, IL-6, IL-18, MCP-1, MCP-4, MDC, MIP-1-a, MIP-1-b, MIP-2-a, TARC, TGF-a, TNF-a, TRAIL-R1, TSLP, VEGF-A, IP-10 and IL-8. Infections were completed with IBV working stocks at a low MOI (0.01) on hNECs. The basolateral media was changed at time zero of the infection. Basolateral samples were collected from the infection at 48 hpi and 96 hpi. Uninfected mock basolateral media was collected at identical time points. Samples were run the custom Procartaplex assay plates with appropriate standards and controls according to manufacturer instructions. Plate was run on a Luminex MagPix System. Raw data was exported and analyzed using ThermoFisher Procartaplex Software and visualized by log fold change to mock in R v4.1.1 using custom scripts.

**Table 7:** MagPix Custom Panel.

| Catalog Number | Name | Lot Number/Bead Lot Number |
| --- | --- | --- |
| PPX-22-MXAACRC | Human Procartaplex Mix and<br>Match 22-Plex | 307393-000 / 305694-000 |

### RNAseq and Analysis

Total RNA at 48 hpi was extracted and purified from hNECs using Trizol reagent (Invitrogen Catalog # 15596018) and the PureLink RNA Mini kit, including on-column DNAse treatment (Invitrogen/ThermoFisher). Quantitation of Total RNA was performed with the Qubit BR RNA Assay kit and Qubit Flex Fluorometer (Invitrogen/ThermoFisher), and quality assessment performed by RNA ScreenTape analysis on an Agilent TapeStation 2200. Unique Dual-index Barcoded libraries for RNA-Seq were prepared from 100ng Total RNA using the Universal Plus Total RNA-Seq with NuQuant Library kit (Tecan Genomics), according to manufacturer’s recommended protocol. Library amplification was performed for 16 cycles, as optimized by qPCR. Quality of libraries was assessed by High Sensitivity DNA Lab Chips on an Agilent BioAnalyzer 2100. Quantitation was performed with NuQuant reagent, and confirmed by Qubit High Sensitivity DNA assay, on Qubit 4 and Qubit Flex Fluorometers (Invitrogen/ThermoFisher). Libraries were diluted, and equimolar pools prepared, according to manufacturer’s protocol for appropriate sequencer. An Illumina iSeq Sequencer with iSeq100 i1 reagent V2 300 cycle kit was used for final quality assessment of the library pool. For deep RNA sequencing, a 200 cycle (2×100bp) Illumina NovaSeq S2 run was performed at Johns Hopkins Genomics, Genetic Resources Core Facility, RRID:SCR_018669. Unaligned FASTQ files and .bam files are available under NCBI BioProject: PRJNA996592.

### Sequencing Analysis

Raw iSeq and NovaSeq FASTQ files were uploaded to the Partek Server and analysis with Partek Flow® (Version 10.0) NGS software, with RNA Toolkit, was performed as follows: pre-alignment QA/QC; trimming of reads; alignment to hg38 Reference Index using STAR 2.7.8a; post- alignment QA/QC; quantification of gene counts to annotation model (Partek E/M, Ensembl Transcript Release 103). Gene counts matrices were exported from the Partek server and further analysis in R v4.1.1. Gene counts were log transformed for normalized analysis using base R rlog. Principle component analy sis (PCA) of normalized gene counts was performed using the plotPCA function from DESeq2 [25]. Differentially Expressed genes were determined using DESeq2 with alpha set to 0.05 for adjusted p-value (padj) thresholding. A gene was differentially expressed at a padj ≤0.05. DEGs were then summarized by a log2fold change of ≥1.5. Gene set enrichment analysis was performed as using gprofiler [26].

**Table 8:** Interferon Gene RT-qPCR – RT qPCR primers.

| Target | Assay | Assay ID |
| --- | --- | --- |
| IFITM1 | ThermoFisher TaqMan | Hs00705137_s1 |
| ZPB1 | ThermoFisher TaqMan | Hs01679797_gH |
| OASL | ThermoFisher TaqMan | Hs00984387_m1 |

Interferon stimulated gene targets identified by RNAseq were performed. Viral RNA was isolated using Qiagen QIAamp Viral RNA Mini Kit per the manufacturer’s protocol. 140µl of nasopharyngeal swab sample was used for each extraction. The concentration of extracted vRNA was measured by NanoDrop and 2µl of vRNA were input into RT-qPCR reactions. Premixed TaqMan primers and probes were run in separate reactions for targets IFITM1, ZPB1, and OASL (Table 1) using TaqPath 1-Step RT-qPCR Master Mix, CG (Thermofisher Cat# A15299). Standard thermal cycling conditions were performed on a QuantStudio 5 Instrument as follows: 1 cycle of Reverse transcription at 50°C for 15 minutes, 1 cycle of Polymerase activation at 90°C for 2 minutes, and 40 cycles of 95°C for 3 seconds to 60°C for 30 seconds of amplification. Analysis was performed using QuantStudio design analysis software v1.3 and statistical significance between groups was tested using 2-way ANOVA in Graphpad Prism.

### Western Blots

Low MOI infections (MOI = 0.01) were performed in hNECs. Cells were extracted using TrypLE (Thermofisher Scientific Cat no. 12604013) and lysed using RIPA buffer (Thermofisher Cat no. 89900). Sample lysates were stained with antibodies raised in rabbit against IFIT2 (Protein Tech 12604-1-AP) at 2ug/ml or IFIT3 (Protein Tech 5201-1-AP) at 1ug/ml. Beta Tubulin-IV raised in mouse was used at a concentration of 1ug/ml as a control. Secondary staining was performed using anti-rabbit AF647 raised in goat and anti-mouse AF488 raised in goat for the IFIT and Beta Tubulin-IV primary antibodies, respectively.

### Data Availability

All RNAseq raw sequence files, .bam files and sample information have been deposited at NCBI Sequence Read Archive, NCBI BioProject: PRJNA996592. All scripts used for RNAseq analysis are available at https://github.com/Pekosz-Lab/IBV_transcriptomics_2023. Raw data used in this manuscript can be obtained through the Johns Hopkins Data Repository at doi XXXXXXXXXX. All genome sequences and associated metadata in this dataset are in the GISAID EpiFlu database. To view the contributors of each individual sequence with details such as Accession Number, Virus name, Collection date, Originating Lab, Submitting Lab and the list of Authors, please refer to Supplementary Table 4.

## Results

### 3.1. Maryland Influenza B Clinical Frequency and Hemagglutinin (HA) phylogenetic assessment

Clinical prevalence of IBV in the United States from 2015-2020 showed prevalence of the B/Yamagata lineage until the 2019-2020 season when the B/Victoria lineage dominated (Figure 1A). Three clinical isolates from each IBV lineage were chosen for further characterization as labeled in Figure 1B. In order to determine relative genetic distance of in- fluenza viruses used in this study to historical and circulating viruses, we performed phylogenetic analysis of the hemagglutinin (HA) using genetically representative sequences obtained from GISAID between the years of 2009 and 2023. A total of 285 complete IBV HA sequences were obtained with accession numbers available in Supplementary Table 4. Maximum-likelihood phylogenetic tree construction of GISAID sequences and HA segments of clinical so- lates revealed distinct clustering by lineage and clade (Figure 1B). B/Victoria used in this study isolated in 2017, B/Baltimore/R0122/2017 and B/Baltimore/R0001/2017, belong to the V1A clade. B/Baltimore/R0696/2020 isolated in 2020 belongs to V1A.3a. All B/Yamagata isolates, B/Baltimore/0300/2018, B/Baltimore/R0250/2018 and B/Baltimore/R0337/2018, belong to the Y3 clade.

**Figure 1.**
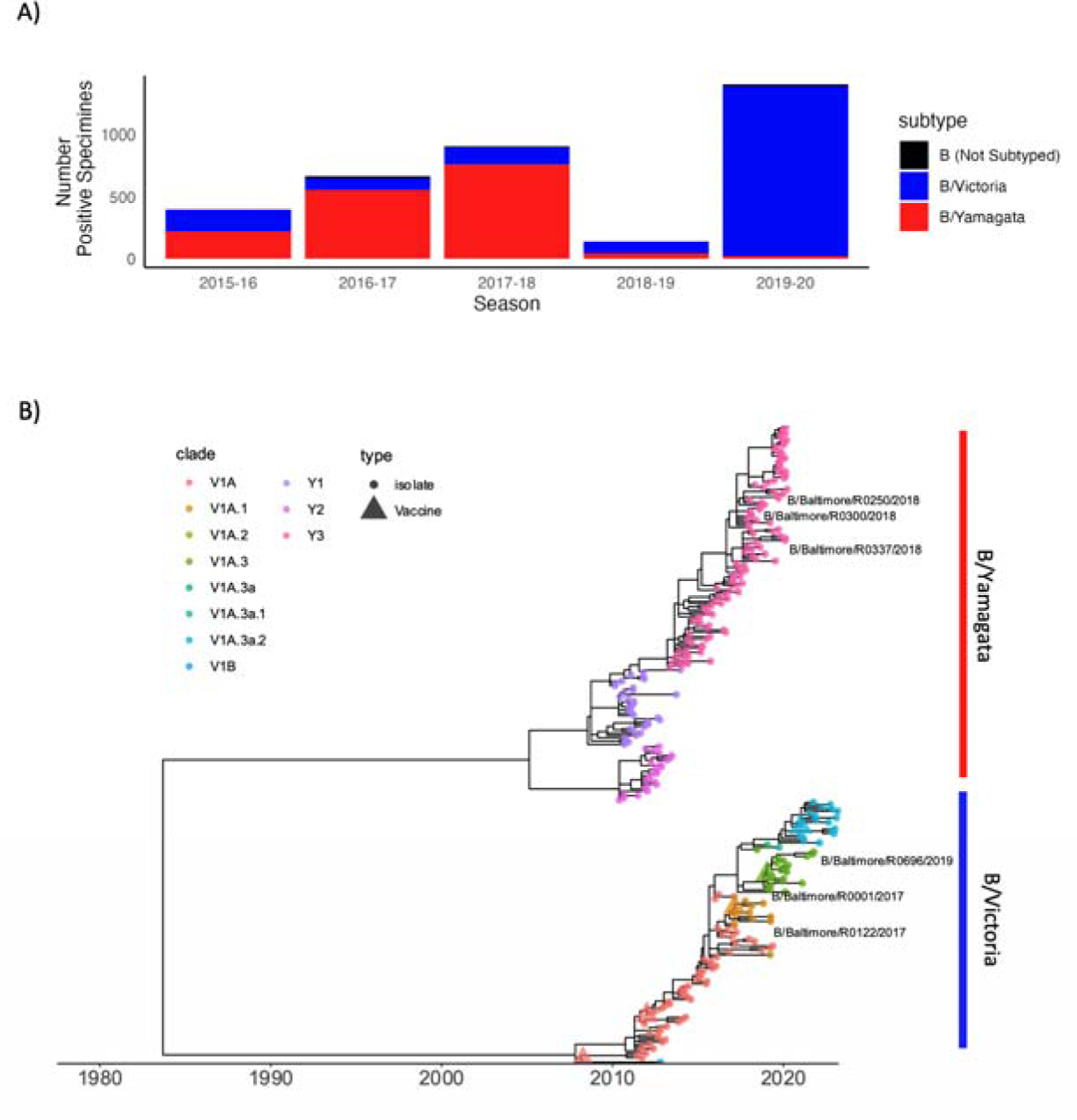
Influenza B clinical incidence and HA phylogenetics. **A**) Number of Influenza B positive clinical specimens as reported by the Centers for Disease Control and Prevention in the State of Marydland from the ‘WHO NREVSS Public Health Labs’ FluView dataset. Data are summarized by lineage and influenza season. **B**) Time-scaled phylogenetic tree of Influenza tree of representative HA sequences (n=282) isolated between 2009-2023. Branch tips are colored by lineage with corresponding vaccine strains defined by tip shape. Isolates collected from the Johns Hopkins Hospital network for subsequent characterization in this study are labeled by isolate ID.

### 3.2 B/Yamagata vaccines induce higher mean post vaccination titers compared to B/Victoria vaccines

A comparison of post influenza vaccination neutralizing antibody titers using serum collected from influenza-immunized healthcare workers at Johns Hopkins University during the 2019-2020 season, was performed to assess how vaccine induced immunity recognized the circulating IBV strains. All participants received quadrivalent inacti- vated vaccine. Neutralizing antibody titers were compared pre and post vaccination to the vaccine strains and the IBV clinical isolates representing the dominant circulating strains. For B/Yamagata that was clade 3 (B/Baltimore/R0250/2018) which is in the same subclade as the vaccine (B/Phuket/3073/2013). For B/Victoria there was an antigenic drift that season and therefore the dominant circulating strain was clade V1A.3 (B/Baltimore/R0696/2019) compared to the vaccine strain, V1A.1 (B/Colorado/06/2017).

Post vaccination neutralizing antibody titers were significantly higher against all viruses tested (Figure 2A). The post vaccination mean neutralizing antibody titers were higher for the B/Yamagata vaccine component compared to the B/Victoria component and titers against the circulating IBV strains were lower compared to the vaccine strain from the same lineage (Figure 2A). Post vaccination titers were higher for the vaccine strains when compared to the circulating viruses in the same lineage. Seroconversion rates, defined as a greater than four-fold increase between pre and post vaccination serum, were slightly higher for the vaccine strains compared to the circulating viruses but these differences did not reach statistical significance (Figure 2B). Together, the data indicate this population had a strong response to influenza vaccination but that the vaccine induced antibodies recognized the vaccine strains better than the circulating strains.

**Figure 2:**
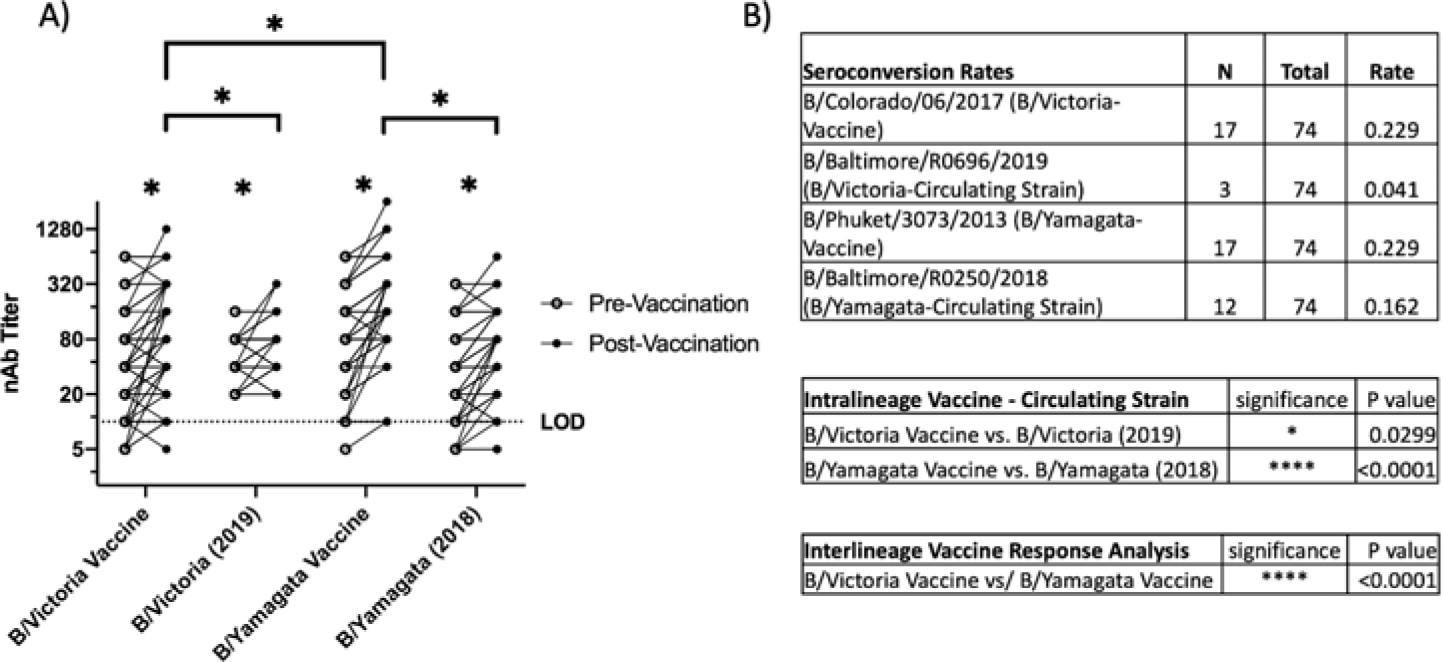
Neutralizing antibody responses to vaccination. A) Neutralizing antibody titers to pre and post vaccination serum. Pre and post vaccination serum was used to compare neutralization of B/Yamagata and B/Victoria vaccine strains (B/Phuket/3073/2013 and B/Colorado/06/2017) as well as B/Yamagata and B/Victoria circulating strains for the year 2019-2020 (B/Baltimore/R0250/2018 and B/Baltimore/R0696/2019). B) Mean difference was calculated, and Sidak’s multiple comparisons test was used to assess signifi- cant differences. Statistical differences set at p ≤0.05 and indicated by *.

### 3.3 Viral fitness is similar between B/Victoria and B/Yamagata viruses isolated between 2016-2019

The serological data suggested that both IBV lineages could evade vaccine induced immunity to a similar level, sug- gesting that lineage specific differences in virus replication kinetics might help explain the dominance of the B/Yamagata lineage in the 2016-17 and 2017-18 influenza seasons (Figure 1A). Using isolates obtained from influenza surveillance efforts in Baltimore, Maryland, we chose to use low MOI infections and comparison of infectious virus production as a measure of in vitro viral fitness. Low MOI infections of the two lineages of IBV showed similar onset of infection as well as peak infectious virus titer in human nasal epithelial cells (hNECs) (Figure 3A).

**Figure 3:**
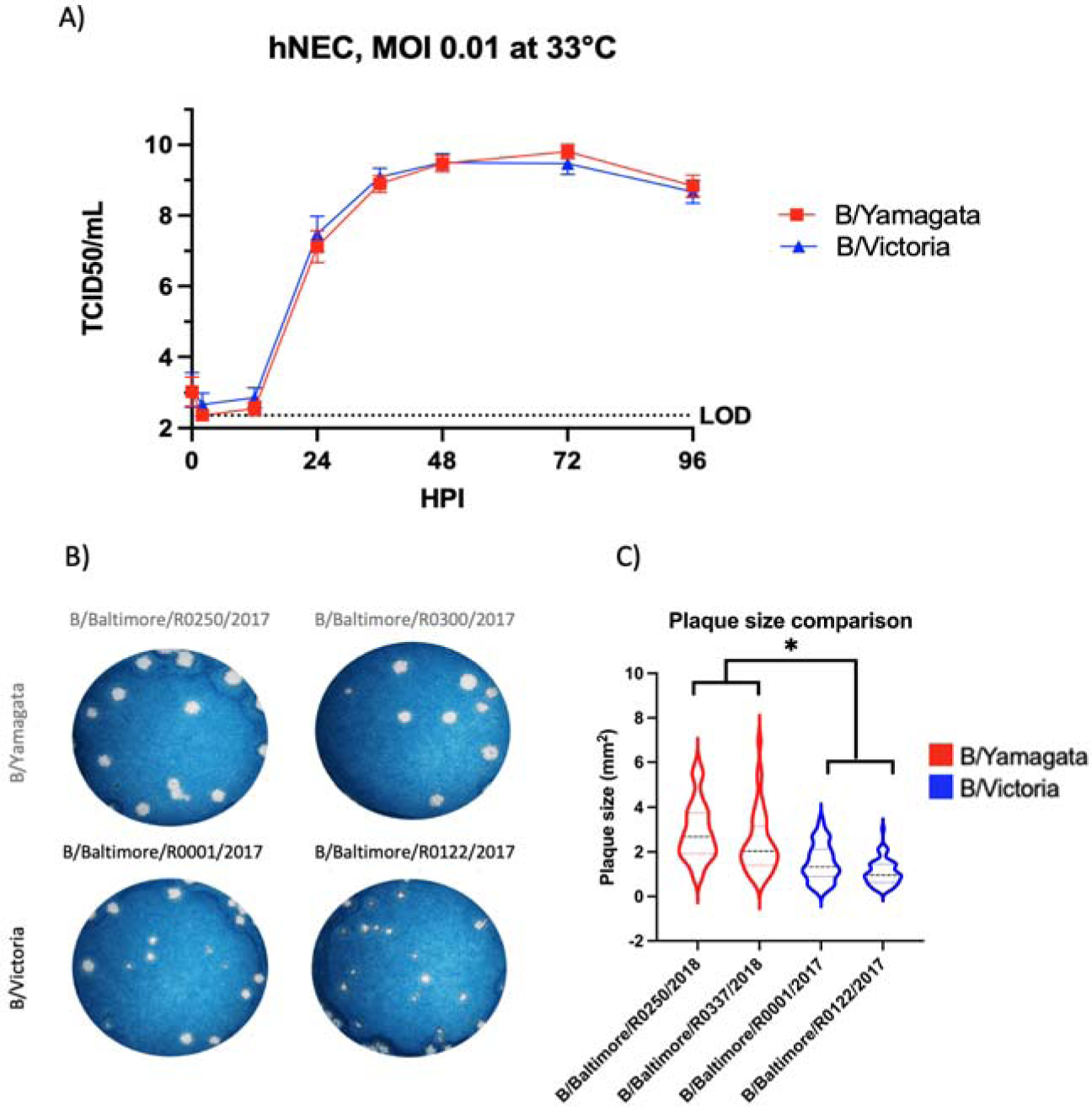
Characterization of Viral fitness between B/Yamagata and B/Victoria viruses. A) Viral fitness assessment on hNEC revealed no significance difference of infectious virus production out to 96 hours post infection of Baltimore clinical isolates. B/Yamagata curves consists of data from B/Baltimore/R0250/2018 and B/Baltimore/R0337/2018 clinical isolates. B/Victoria curves consist of data from B/Baltimore/R0001/2017 and B/Baltimore/R0122/2017. Growth curves completed in two independent experiments, each experiment consisted of three replicate wells .Statistical significance of growth curves calculated on GraphPad Prism using a two-way ANOVA and Tukey’s multiple comparisons B) Representative wells shown comparing plaque size from two independent experiments of the two lineages of IBV on MDCK cells. All plaques in experiment wells were counted for analys s. C) Comparison of plaque sizes formed by two clinical isolates from each IBV lineage. Graphed plaque areas are the combination of two independent experiments. Images of plaques were analyzed with ImageJ.

Plaque size and morphology is a common technique used to evaluate viral phenotype. Plaque formation allows an assessment of cell to cell spread in the MDCK cell model. We directly compared IBV plaque formation between the B/Victoria and B/Yamagata lineage viruses. B/Victoria viruses consistently show smaller plaque size (Figure 3B-C) compared to B/Yamagata viruses (p < 0.0001).

### 3.5 Influenza B viruses infect multiple types in the nasal respiratory epithelium however predominate in the ciliated cells

Bui et al used immunohistochemistry (IHC) to show that IBV clinical isolates from both lineages infect multiple cell types in the bronchial respiratory epithelium. We sought to define cell tropism in our hNEC model. Using flow cytometry, we can gain a quantitative view of IBV infection over the initial five days of infection. IBV infected hNECs were isolated by first gating out debris, multiplets and dead cells followed by gating of IBV HA positive cells (Figure 4A). We used intracellular markers commonly used to describe various cell types in the respiratory epithelium. Beta-Tubulin-IV (BT-IV) was used to define mature ciliated cells [27], [28].The hNEC cultures showed three populat ons of BT-IV staining (Supplemental Figure S1A). Total populations include a negative staining population, an intermedi- ate population and a BT-IV positive population. When fluorescence was plotted against side scatter area (SSC-A) a pass both goblet cells as well as ciliated cells that produce mucus as has been well defined in the literature (Supplemental Figure 1A) [30], [31]. NGFR (CD271) was used as a marker for respiratory basal cells.28 NGFR staining in our hNEC cultures did not show any significant co-staining with the brightest BT-IV population defined as ciliated cells or MUC5AC containing cells consistent with expected protein production in these developing cell types (Supplemental Figure 1B,D). Once these cell types were identified, we used this method to identify quantitative IBV cell tropism of our infected hNEC cultures. At peak infectious virus production, 72 hpi, the majority of IBV infected cells were ciliated cells (B/Victoria 54.77% and B/Yamagata 52.18%, p=0.599) followed by mucus producing cells (B/Victoria 35.45%, B/Yamagata 39.04, p=0.155) and basal cells (B/Victoria 4.81, B/Yamagata 11.06, p=0.0045) (Figure 4B). There was no significant variation between lineages in the ciliated cell or mucus producing cell populations. B/Yamagata viruses infected significantly higher NGFR + basal cells compared to B/Victoria (Figure 4B).

**Figure 4:**
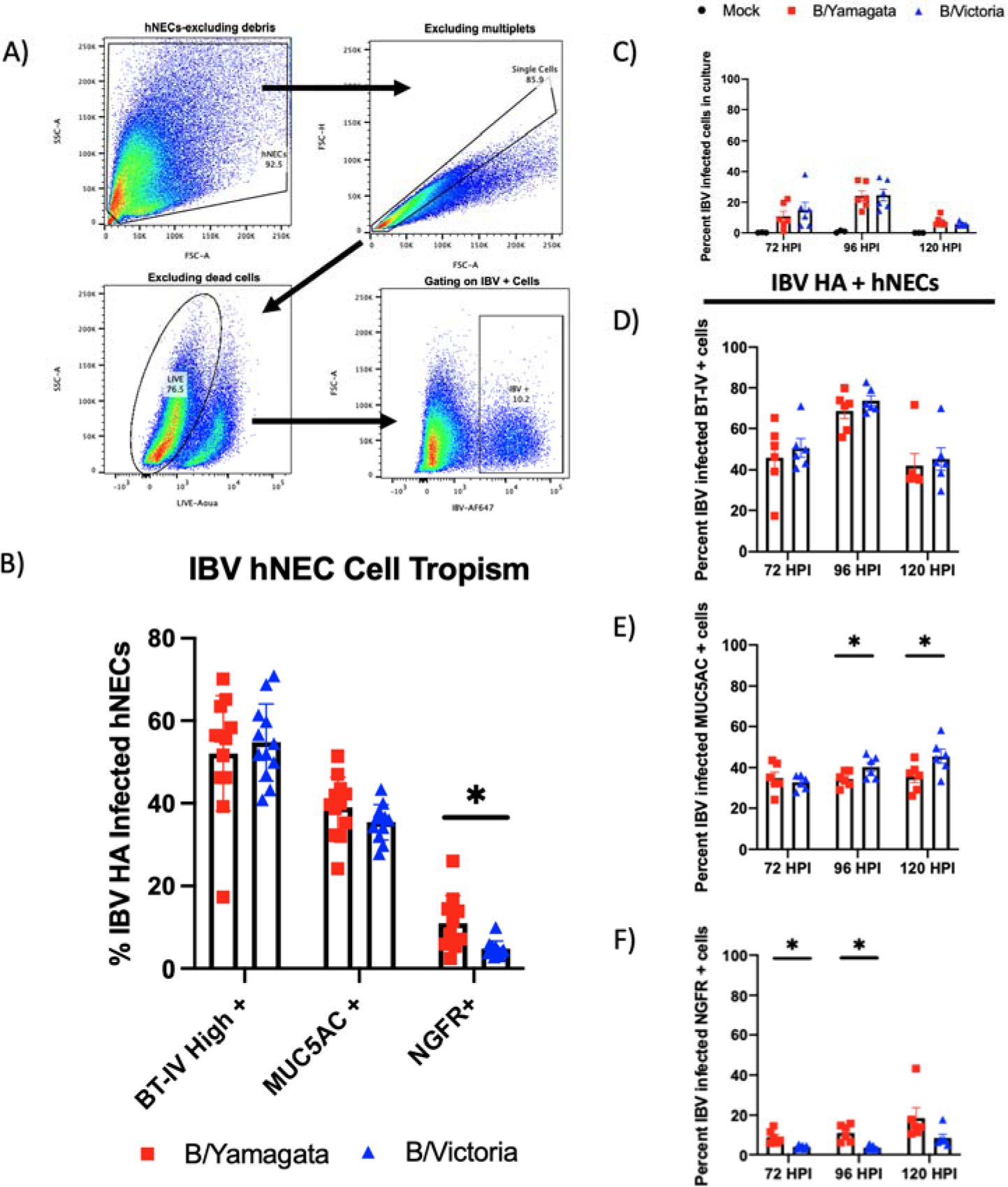
Cell tropism assessment of IBV infection of human nasal cells. A) Gating strategy of assessing IBV infected hNECs using flow cytometry. Cells were gated by removing debris, single cells, live cells and IBV HA +. B) Additional marker of human nasal cells were used to assess cell tropism including BT-IV as a marker of mature ciliated cells, MUC5AC as a marker of mucin producing cells and NGFR(CD271) as a marker of respiratory basal cells. Statistical Analysis: Percentages of IBV infected hNECs were statis- tically compared between two groups using multiple t –tests. Data represents 72 hour infection data from two independent runs ed hNECs were identified using HA monoclonal antibodies for IBV HA. The specific cell types infected over the course of infection were identified using antibodies specific for the same cellular targets as shown in panel B. (C) Percent of IBV infected cells over infection course. (D) Percent of IBV infected ciliated cells infected through 120 hours. (E) Percent of IBV infected MUC5AC + cells through 120 hours. (F) Percent of IBV infected NGFR+ cells through 120 hours. Cell populations were identified with the following gating strategy: excluding debris, single cells, Live cells, IBV + cells, specific cellular stain as described. Three replicate wells were completed for this experiment. Data from two clinical isolates included per lineage. Statistical Analysis: Percentages of IBV infected hNECs were statistically compared between two groups using multiple t –tests.

### 3.6 IBV infected ciliated cells peak at 96 hpi where infected basal and mucus producing cells continue to increase through the course of infection

We know from our replication kinetics experiments that infectious virion production of influenza B viruses peaks at 72 hpi (Figure 2A). In contrast to our infectious virion production experiments, when we evaluated the number of infected cells over time we saw an increase that peaked at 96 hpi prior to decrease (Figure 4C), with approximately 20% of the total cells infected at that time. The infected ciliated cells drive this pattern as the most infected cell type with peak number of infected cells at 96 hpi (Figure 4D). In contrast to the ciliated cells, IBV infected MUC5AC producing cells continue to increase throughout the course of acute infection (Figure 4E). This pattern was consistent with what was noted in infected basal cells. The previously noted pattern of increased IBV basal cell infection in B/Yamagata infected cultures was again seen at 96 hpi but was not statistically significant at 120 hpi (Figure 4F).

### 3.7 Pro-inflammatory cytokine and chemokine production induced by Influenza B Infection was predominantly defined by IL-6, G-CSF, MCP-1 and TGF-a

To expand the understanding of innate immune responses in influenza B virus infection using protein level immunoassays, a custom Luminex panel was designed to detect commonly upregulated cytokines and chemokines in response to infection with influenza and other respiratory viruses [32]. Cytokine and chemokine production was evaluated both at 48 hpi and 96 hpi. Cytokine production is about 4-fold higher at 96 hpi compared with the 48-hour time point (Figure 5A-B). Although significantly greater production, relationships of cytokines and chemokines produced remained relatively similar between time points. CXCL-10, IL-6, G-CSF, MCP-1, TGF-a, TNF-a, MIP-2-A, BAFF, MDC, TRAIL-R2 and MIP-1-A, all had at least 2-fold change from mock infected wells although many proteins in the panel showed some degree of upregulation post infection (Figure 5A-B). CXCL-10 had the highest degree of upregulation between 30-50-fold mock at 96 hpi (data not on graph given scale difference). There was no significant difference between cytokine production in response to infection when comparing B/Yamagata and B/Victoria (Supplemental Table 1).

**Figure 5:**
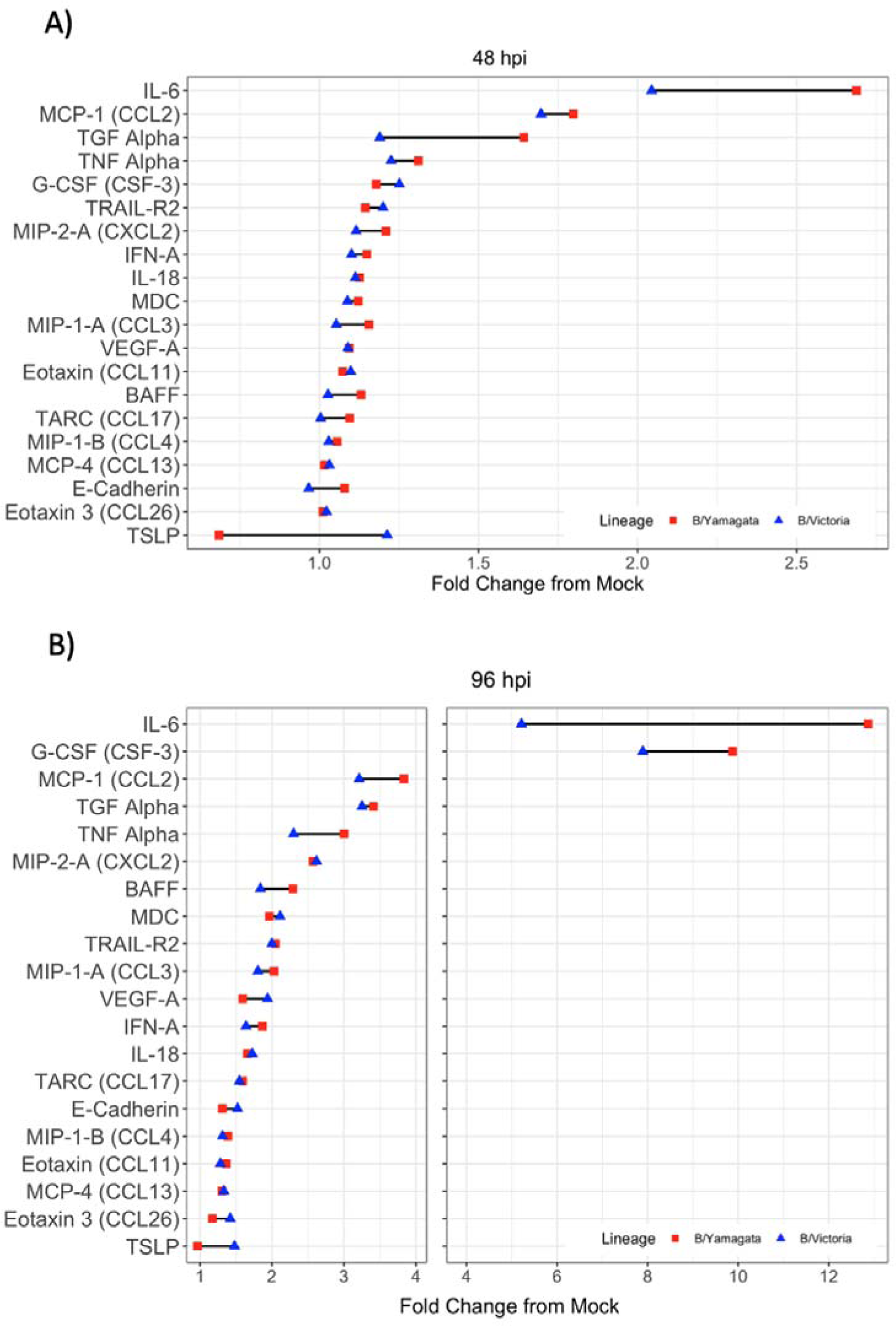
Cytokine expression profiles in the basolateral supernatant of IBV infected hNEC cultures. General profiles of inflammatory signature of IBV infected hNECs is shown comparing infections with B/Yamagata and B/Victoria clinical isolates. Data represents two independent experiments. For each experiment, three replicate wells were ana- lyzed. Each IBV lineage combines data from two clinical isolates. There was no statistically significant differences between cytokine profile following infection between B/Yamagata and B/Victoria.

### 3.8 RNAseq of B/Victoria and B/Yamagata Infected hNECs

To expand on post infection innate responses, bulk RNAseq was performed to determine the gene expression profiles Jo4_337_1_S36 (Yamagata_337 Replicate 1) was excluded from analysis as 75% of reads belonged to non-unique sin- gletons which mapped to ribosomal RNA at an average coverage depth of 275 (Supplementary Table 5). The remain- ing samples averaged a sequencing coverage depth of 71.8 across treatment groups. All included samples resulted in high mapping percent alignment to human genome hg38 at an average of 92.2% (Supplementary Table 5). Hierarchical clustering of rlog transformed count tables indicated strong clustering between samples at the lineage and virus-specific level (Supplementary Figure 2A). Principle Component Analysis (PCA) of rlog transformed read counts resulted in strong separation by PC2 between IBV lineages and mock treatment of hNECs with the represented vari- ance of PC1 and PC2 to be 58% and 32%, respectively (Supplementary Figure 2B).

### 3.9 Differentially Expressed Gene (DEG) analysis identifies strong upregulation of Type I and III interferon stimulated gene families

As all viruses chosen within each lineage belong to identical clades, we chose to focus our DEG analysis by lineage to identify major trends across B/Victoria and B/Yamagata infection. We compared DEGs post B/Victoria and B/Yamagata infection against mock using DEseq2. Assessment of global transcription across treatment groups revealed distinct profiles in the top 112 DEGs with an adjusted p value ≤ 0.05 without fold change filtering. Z-scores from rlog-normalized gene counts represent relative expression (Figure 6A). Hierarchal clustering by genes and individual virus expression patterns resulted in distinct clustering by both virus and lineage across all DEGs. Further filtering by both adjusted p-value ≤ 0.05 and a log2 fold change ≥1.5 revealed that many top upregulated genes belong to the interferon stimulated gene family and are more highly expressed in B/Victoria infected hNECs (Figure 6B). Differentially expressed genes were further assessed by a log2 fold change ≥ 0.8 and padj ≤0.05. A total of 115 genes were upregulated in B/Victoria infected hNECs as compared to mock as to whereas only 54 upregulated DEGs were identified in B/Yamagata infections at this threshold (Figure 8A). 53 upregulated genes were shared between the two lineages with 40 of these genes expressed greater in B/Victoria infections (Figure 6B). CXCL10 and ZPB1 were among the top 5 DEGs in both lineage infections. The remaining genes comprised of interferon and interferon stimulated genes (ISGs) in both B/Victoria and B/Yamagata. IFN-l (IFNL1, IFNL2, IFNL3) were among the top upregulated genes in both lineages along with ISG belonging to the IFIT, IFITM and OAS family including IFI27, IFTM1, IFI6, OAS2 and OAS3 (Figure 8B-F) all of which were expressed greater in B/Victoria. Shared genes upregulated with a higher log2 fold change in B/Victoria infection included known anti-viral proteins IFIT1-3, IFITM1, MX2, IFI44L and ISG15 along with mitochondrial-related gene CMPK2. Implicated ISG15 E3 ligase, HERC6 was also highly upregulated in B/Victoria along with HERC5 and TRIM31 [33]. 49 out of 115 genes were uniquely upregulated in B/Victoria compared to all treatment groups including CXCL11, IFITM2, EIF2AK2, HLA-B along with many long non-coding RNAs. Only one DEG was identified to be unique in B/Yamagata infection, ENSG00000204745, which is an anaphase promoting complex subunit (ANAPC1) pseudogene. Using adjusted p-value, no downregulated DEGs were shared between B/Victoria and B/Yamagata infected hNECs (Figure 7C). Few downregulated genes were identified in B/Victoria infections. COL12A1, a collagen type XIII factor, was downregulated in both B/Victoria infections relative to mock along with ENSG00000274944, a novel unnamed protein. No DEGs were identified as being unique in B/Yamagata infection.

**Figure 6:**
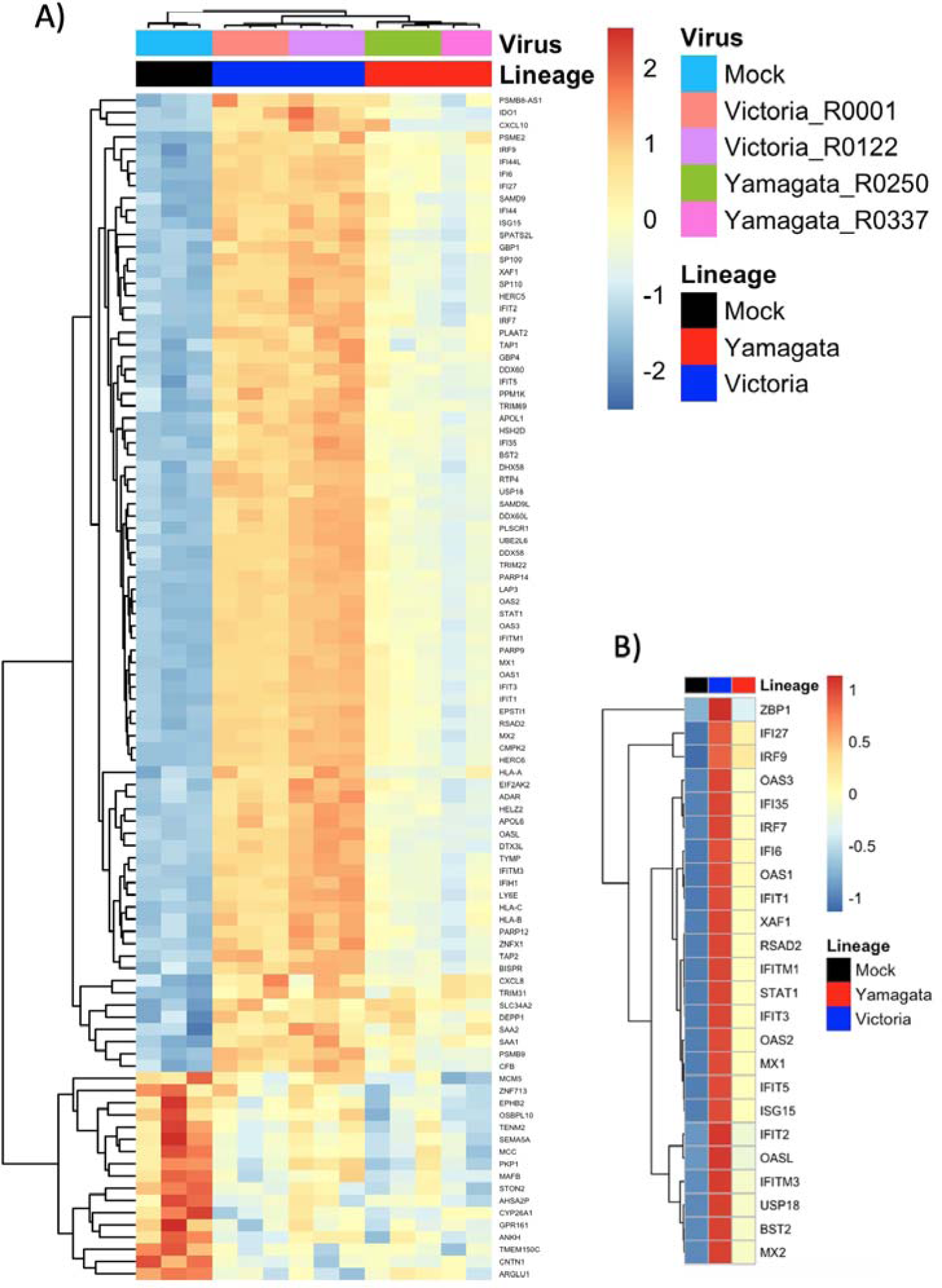
hNEC Transcriptional response to B/Victoria and B/Yamagata infection (A) Heatmap of DEGs with a padj ≤ 0.05 in hNECs (n=122). (B) Subset the complete heatmap of annotated by the top 24 differentially expressed genes summarized by B/Yamagata and B/Victoria lineages.

**Figure 7:**
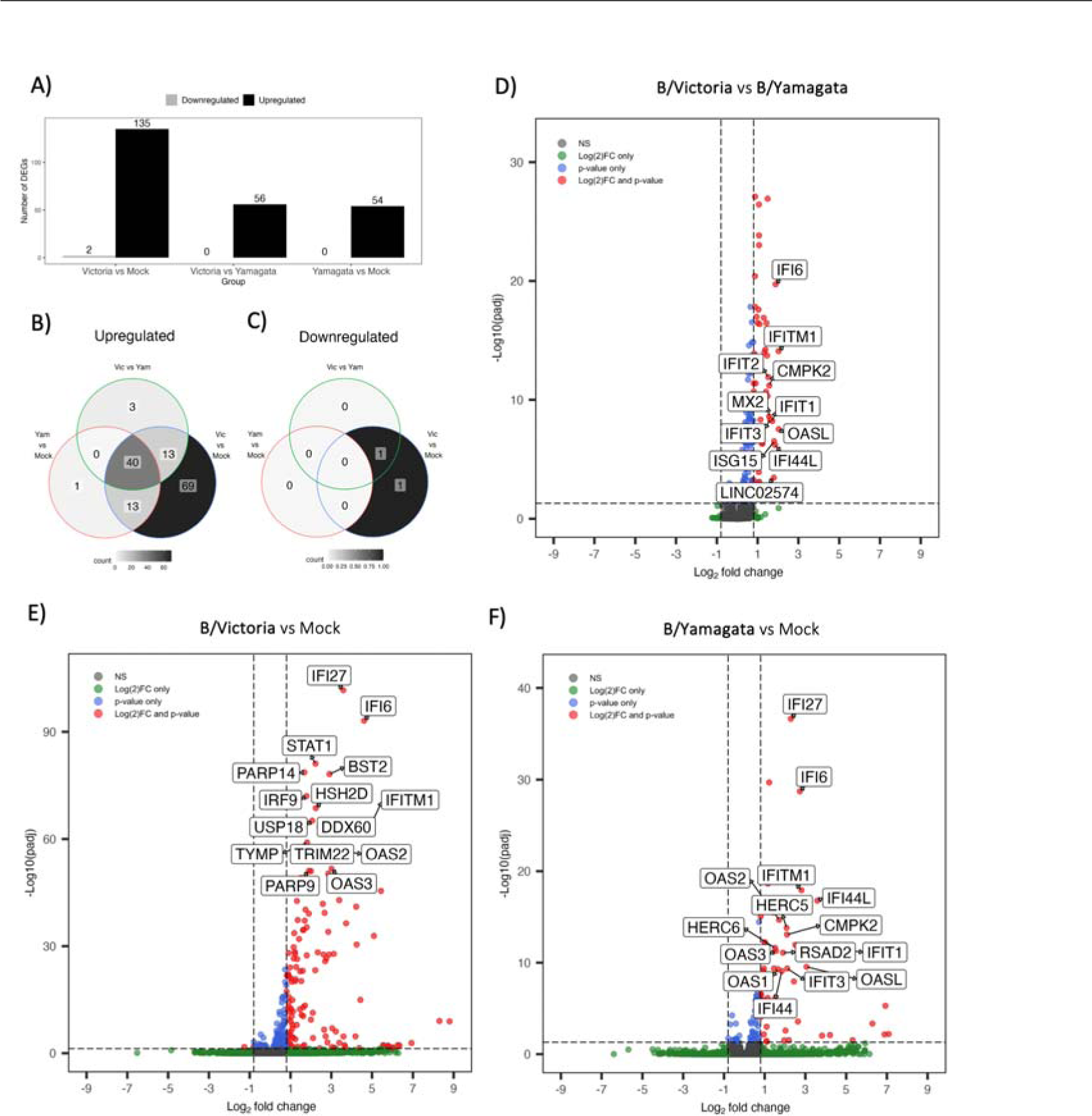
Differentially expressed genes between B/Victoria and B/Yamagata infected hNECs. Total Differentially expressed genes (DEGs) at threshold cutoffs of a log2 fold change ≥0.8 and padj ≤0.05. (A) Total up and downregulated genes at threshold cuto fs representative of all DESeq2 design comparisons. Venn diagrams for each design comparison separated by up (B) and down (C) regulated genes. Volcano plots of DEGs at threshold for B/Victoria vs B/Yamagata (D) B/Victoria vs Mock (E) and Yamagata vs Mock (F).

**Figure 8:**
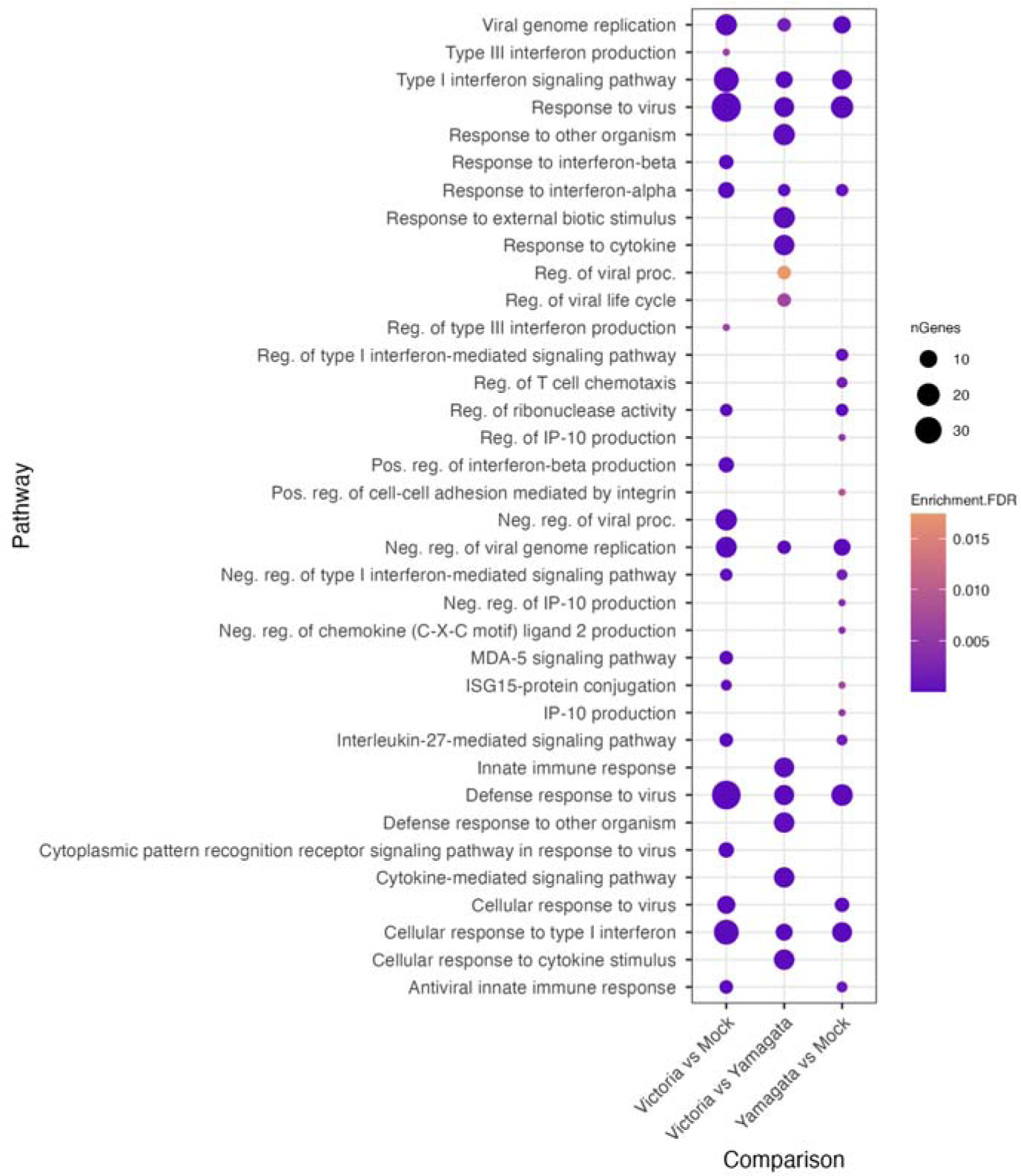
Pathways differentially regulated during B/Victoria and B/Yamgata infection. Pathway enrichment plot representing the top enriched pathways across all lineage comparison using DEseq2 with threshold cutoffs of padj ≤0.05 and log2 fold change ≥0.8 using gprofiler.

### 3.10 Gene Ontology Analysis

To further identify differentially expressed gene pathways of infected hNECs, we performed gene ontology analysis on identified DEGs. Only upregulated genes were considered. Analysis was performed on each DEG analysis for all DESeq2 comparison groups by lineage. First, Gene Set Enrichment analysis (GSEA) was performed on DESeq2 output across all comparison groups: B/Victoria vs Mock, B/Yamagata vs Mock and B/Victoria vs B/Yamagata. We focused on a broader range of genes filtering for a log2 fold change ≥ 0.8 and p value ≤ 0.5 to evaluate pathways. No unique pathway differences were identified between lineage treatment groups and belonged to the antiviral response systems, cytokine signaling type I, type III interferon pathways, and Negative regulation of viral replication or processes (Figure 8). B/Victoria vs Mock groups consistently identified a higher number of genes in each other shared pathways belonging to Cellular response to type I interferon and, Defense Response to Virus, Type I interferon signaling pathway and negative regulation of viral processes driven by the unique subset of genes upregulated by B/Victoria infection.

### 3.11 mRNA and Protein Expression of RNAseq Targets

To validate mRNA differential expression from RNAseq, 48 hpi hNEC infections were repeated as described above. Cell lysates were subjected to RT-qPCR and western blotting for identified upregulated genes in the interferon response gene and antiviral pathways. Total mRNA for ifitm1, zpb1 and oasl were induced after viral infection from both lineages representing results consistent with RNAseq data (Figure 9A-B). For protein production validation, ISGs FIT2 and IFIT3 were selected and quantified with western blot. Induction of IFIT2 and IFIT3 were observed in hNECs across both lineage infection groups with the strongest signal observed in B/Yamagata-infected cells for IFIT2 with similar amounts of protein detected across all virus infections for IFIT3 (Figure 9C).

**Figure 9:**
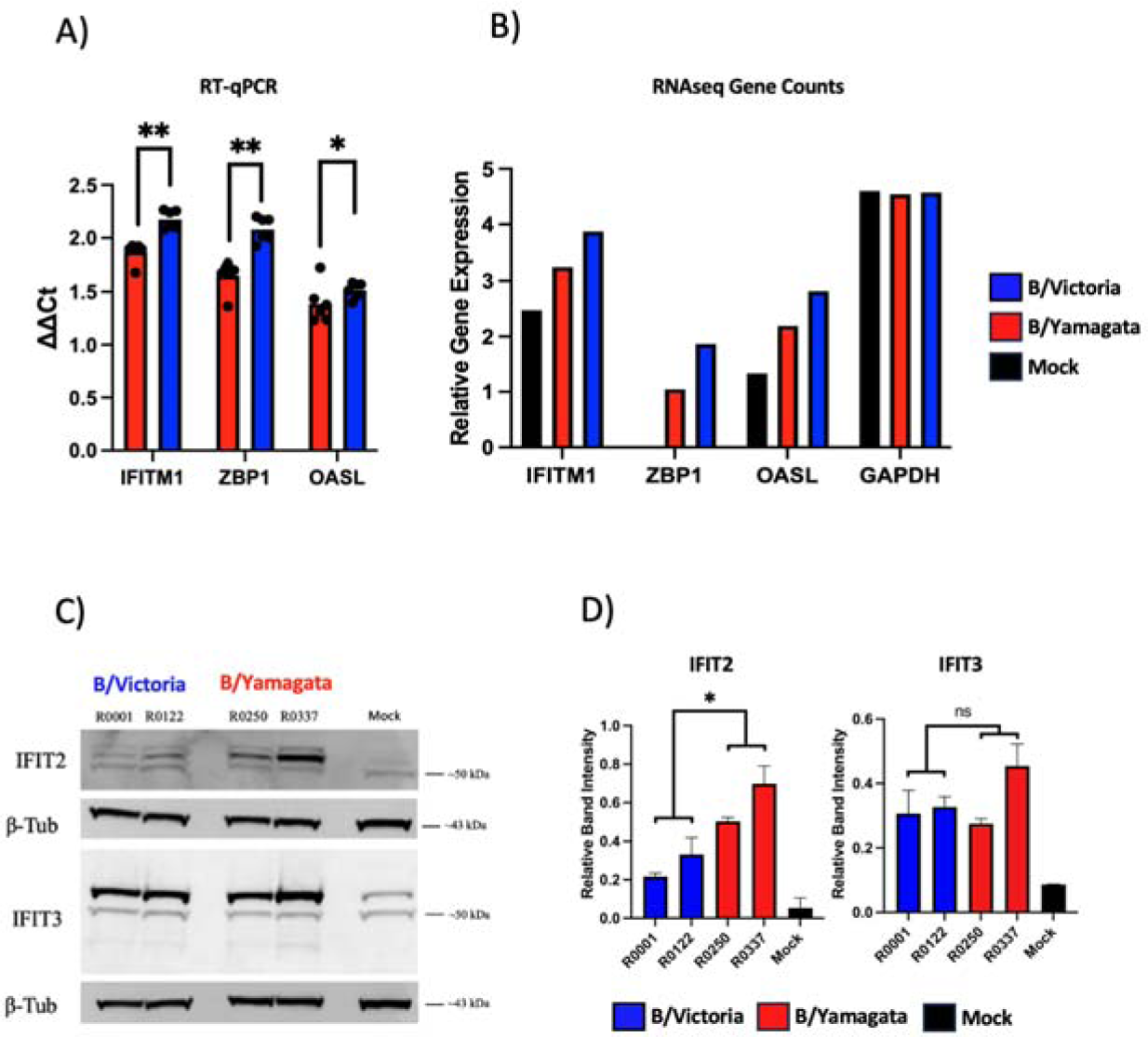
RNAseq validation by RT-qPCR and Western Blot A) RNAseq validating using RT-qPCR of 5 targets: OASL, IFITM1, RBFOX3, USP17L1 and ZBP1. Change is gene expression was calculated using the 2^-(ΔΔCt)^ method and summa- rized by lineage. Comparisons were performed by two-tailed t-test using Graphpad Prism. B) Relative gene expression counts for each validated gene from rlog normalized RNAseq counts. C) Western blot and relative band intensities of (D) IFIT2 and IFIT3 for each virus by lineage from two independent experiments. Statistical significance of normalize band intensities were calculated on GraphPad Prism using a two-way ANOVA and Tukey’s multiple comparisons.

### 3.12 IBV NS1 Sequence Analysis

Given that the pathways which were differentially upregulated between lineages belonged to the interferon and anti- viral related responses, we sought to compare IBV genomes focusing on a known IBV antagonist of these pathways. IBV NS1 is known to be a critical factor in inhibition of antiviral response systems, including but not limited to its ability to bind host proteins such as ISG15 and IFIT2 1,2. To further explore if variation in NS1 might account for lin- eage level immune response differences, we chose to explore phylogenetic differences in B/Yamagata and B/Victoria. To evaluate NS1 protein diversity, 254 NS1 protein sequences were accessed from GISAID belonging to both B/Victoria (n=142) and B/Yamagata (n=108) lineages and aligned to the four viruses used in this study. Phylogenetic analysis of IBV NS1 by maximum likelihood tree construction of the NS1 protein revealed high divergence between the lineages (Figure 12A). Intra-lineage divergence between viruses used in this study show higher distance between the B/Yamagata viruses compared to the B/Victoria. Analysis of all NS1 sequences was used to generate consensus se- quences for alignments to both lineages along with overall amino acid frequency and richness (Figure 12B). Specific amino acids from consensus sequences were unique by lineage. These include a N-terminal asparagine insertion at position 3 observed exclusively in all B/Victoria viruses. The linker region contained additional regions unique by consensus observed at positions 4, 7, 111, 115, 118, 120, 127 and 139. Calculating Shannon entropy by alignment site revealed the highest amount NS1 variation to be at the N terminal alignment positions 3 and 4 and residues within the linker region from 110-140 (Figure 12B). The C-terminal region contained fewer variable residues with a notable K177R B/Victoria to B/Yamagata consensus lineage difference. The remaining entropy belonged to intra-lineage variation.

**Figure 10:**
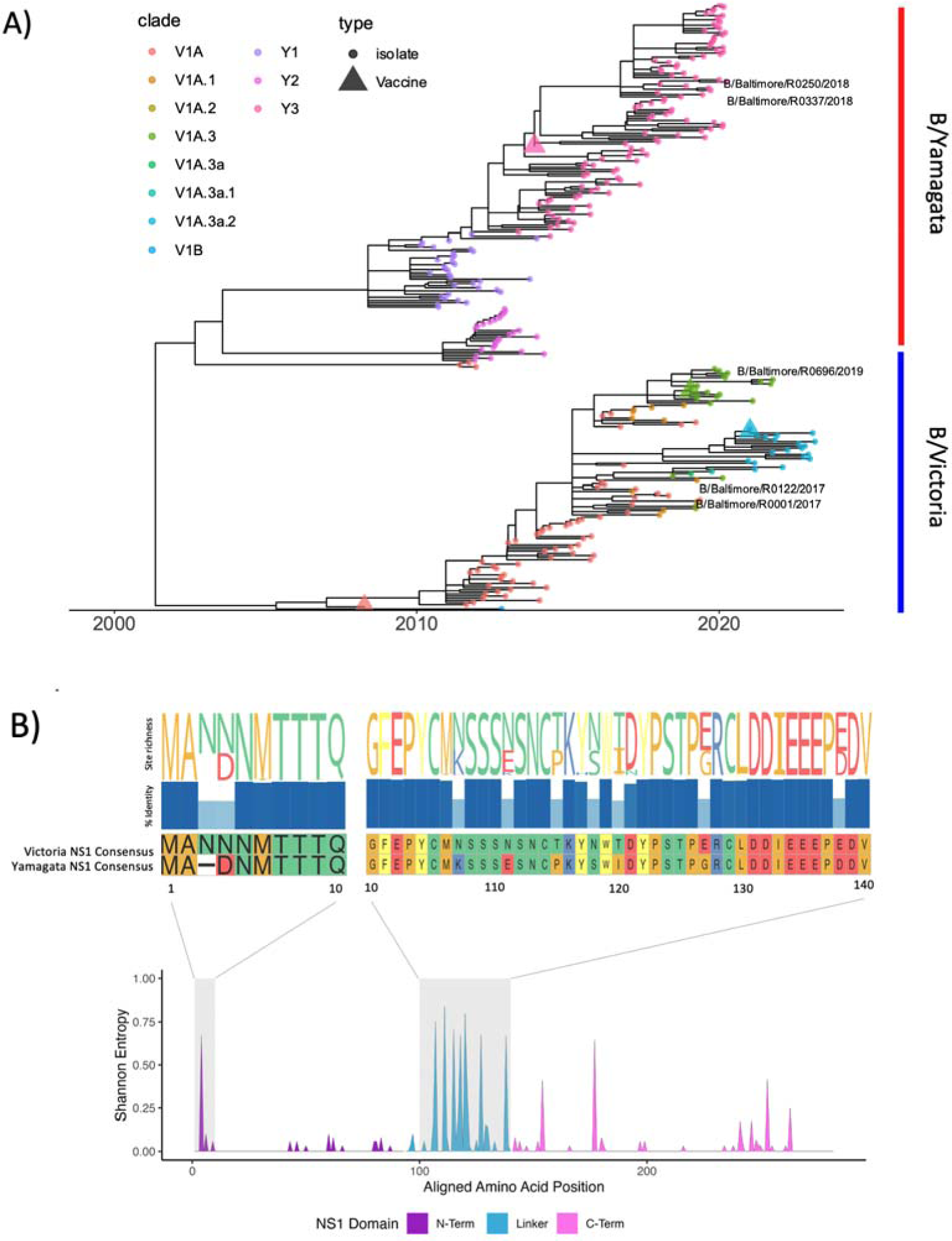
Influenza B NS1 protein has diverged by lineage and HA clade A) Influenza B phylogenetic tree of representative NS1 sequences (<u>n=283</u>) isolated between 2009-2023. NS1 open reading frames were extracted using the NCBI Influenza annotation ool (https://www.ncbi.nlm.nih.gov/genomes/FLU/annotation/) and aligned using *Muscle* (v3.8.31) with the PPP algorithm. Time-scaled maximum-likelihood trees were constructed using *treetime* v0.9.6 and annotated by lineage in R (v4.1.2) using *ggtree* v3.16. B) Amino acid alignments were used to calculate Shannon entropy by site using the *vegan* v2.6-4 and *seqinr* 4.2-23 packages and visualized using R v4.1.1. IBV NS1 domains were annotated according to PDB: 5DIL and BMRB: 25462. Amino acid site richness and percent identity were visualized using *ggmsa* 3.16.

## Discussion

IBV has been given less research attention compared to IAV likely due to smaller proportion of annual infections and decreased pandemic potential due to lack of diverse animal reservoir [34]. IBV historically has accounted for approx- imately one-quarter of the influenza infections and has a major impact on the pediatric population, especially during the northern hemisphere in 2021. B/Yamagata has not been sequenced since March of 2020. Although many hypothesize that this lineage may be extinct, IBV has seen pauses like this in the circulation historically therefore more surveillance is needed to fully assess B/Yamagata disappearance [14]. The primary purpose of the study was to compare the acute respiratory epithelial infection and immune response to IBV infections as a whole and further to compare the two lineages of IBV. We initially hypothesized based on varying epidemiologic patterns and age predilection that these two virus lineages may behave differently in terms of viral replication and immune response. We used pre and post vaccination serum to evaluate antibody production between lineage. We used hNECs to model acute infection and assessed viral fitness, nasal respiratory cell tropism, protein immunoassays and transcriptomics. Additionally, we used MDCK cell models to evaluate plaque size.

Lau et al recently evaluated hemagglutinin inhibition (HAI) titers comparing B/Yamagata and B/Victoria in IBVs isolated from 2009-2014. They found that in adults, B/Yamagata viruses lead to higher antibody responses to vaccination. To evaluate this using a different method, we used serum neutralization antibody assays [35]. Our findings are consistent with Lau et al in a higher mean post vaccination antibody response in B/Yamagata compared to B/Victoria. We hypothesize that this could be due to antigenic imprinting given that B/Yamagata clade 3 has dominated as circulating clade since 2012 whereas recent B/Victoria genetic diversity has given rise to a new dominating clade roughly every two years since 2015. Our findings of differences between vaccine and circulating strain are not surprising given known drift of B/Victoria in the 2019 season and the accumulation of egg adaption mutations at antigenic sites of IBV vaccine strains [36]

In the 2016-2018 seasons, B/Yamagata, clade 3, viruses dominated and B/Victoria V1A.1 circulation was diminished in favor of the alternate lineage (Figure 1A). We assessed viral fitness using replication growth curves between these viruses, finding similar onset of infection, and burst size. We conclude that innate viral replication factors likely do not play a role in the pattern of shifting dominance between these viruses in circulation and rather likely is due to patterns of immune memory in the population as has been studied by several groups [37], [38]. Our plaque evaluation between these lineages showed an increased plaque size with B/Yamagata viruses suggesting that there are some innate viral factors making these lineages distinct.

We set out to understand the initial 5-day course of infection in the nasal respiratory epithelium beyond replication kinetics. Bui et al showed that IBV infects multiple cell types at 37 °C in the bronchial epithelium [39]. We chose to take a quantitative approach of the nasal epithelium using low MOI hNEC infections and analyzing infected cell numbers using flow cytometry. Our data agrees with the Bui et al. and adds to this knowledge by highlighting the changing dynamic of the infected nasal epithelial subsets during acute infection. At the peak of infection (72 hpi), the majority of infected cells are ciliated cells followed by mucus producing cells and then basal cells. There was a statistically significant finding of increased basal cell infection in B/Yamagata compared to B/Victoria, whereas other infected cell type percentages were not different. As we showed a variation in plaque size as well in this lineage we conclude that this may mean that cell to cell spread mechanisms may be different between lineages. This pattern seen in B/Yamagata deserves further study. We showed that although virion production peaks at 72 hpi, number of infected cells peaks at 96 hpi. This pattern is driven by the ciliated cells given highest proportion of infected cells. The mucus producing cells and the basal cell populations follow a different pattern of increasing throughout the course of infection. This is likely secondary to IBV infection mediated cell death of the ciliated cells in early infection leading to increased exposure and therefore vulnerability of other cell types in late infection.

A broadly descriptive phenotype of the immune response to IBV infection, to our knowledge, has not been reported. We chose to evaluate hNEC culture responses to infection using both protein immunoassays as well as bulk RNAseq. Protein immunoassays showed highest epithelial production of CXCL10, IL-6, G-CSF, MCP-1 and TGF-α. A similar study in IAV hNEC infection showed similar upregulation patterns although this evaluation was at the mRNA level [40]. To our knowledge, our study is the first to employ distinct RNAseq characterization of the hNECs infected with IBV at the lineage level. At 48 hpi, we observe a large number of upregulated DEGs belonging to the interferon and antiviral pathways including interferon stimulated genes (ISGs). Upregulation of OAS, IFIT and IFITM gene families in all IBV infection align with previously cited studies performed in immortalized cell lines within in a hNEC culture system. CXCL10 (Supplementary Table 3) was highly expressed across both B/Victoria and B/Yamagata which is consistent with previous clinical reports in IBV host response from nasopharyngeal swabs [41]. While both lineages share many genes belonging to the ISG response, RNAseq reveals that B/Victoria transcriptional responses are higher in both multitude and magnitude of ISGs as well as uniquely upregulating genes such as CXCL11, IFITM2 and STAT2 compared to B/Yamagata infections.

ISGs IFIT2 and IFIT3 were strongly expressed in both lineage infections as verified using western blotting. However, amounts of IFIT2 and IFIT3 appear to be slightly higher in B/Yamagata infections with an exceptionally strong band in B/Baltimore/R0337/2018 infection. The importance of these proteins during influenza infection is well demonstrated with evidence of a dual pro-viral function to bias viral transcript production in both IBV and IAV in vitro infections [2], [7].

ISG15 ubiquitin-like protein is upregulated in both lineage treatments with stronger transcript abundance measured in B/Victoria infected hNECs. Several studies have demonstrated that this protein is important for viral replication control in vitro [42], [43].

It is important to consider that the IBV lineages may have evolved different strategies for immune avoidance. Like IAV, influenzas B encodes for a non-structural protein on segment 8 which primarily acts to counteract innate host immune proteins deemed NS1. Uniquely, the IBV NS1 encodes for extra RNA-binding residues in the C-terminal domain and has demonstrated unique binding specificity to host antiviral proteins such as ISG15 compared to IAV [11], [43]–[45]. More attention is needed to characterize the host response to IBV lineages as B/Victoria NS1 protein evolution is consistent with that of the HA following the emergence of novel lineages since the SARS-CoV-2 pandemic.

Without a known animal reservoir, Influenza B viruses must maintain a balance of fitness and host immunity evasion through variation in the HA and NA segments. Our analysis reveals high diversity and lineage-specific divergence of the IBV NS1 protein which is consistent with HA diversification. We hypothesize that this NS1 diversity contributes to distinct host response in our hNEC model.

We set out to describe in detail the in vitro properties of IBV lineages and the hNEC culture response to infection with different IBV lineages. We conclude that IBV replication peaks around three days on respiratory epithelial cells with no significant differences in fitness between modern clades of B/Victoria and B/Yamagata suggesting immune mechanisms as more likely contributing to shifting seasonal dominance. We conclude that there may be differences in cell to cell spread between B/Victoria and B/Yamagata based on plaque phenotype and basal cell infection frequency. And finally, we describe the transcriptional response seen between infection of B/Victoria and B/Yamagata.

## Supplementary Materials

Supplementary Figure 1. Identification ofIhNECIsubpopulations by flow cytometry. A) FlowJo Plots to show gating of hNEC subpopulations. BT-IV high cells were gated using SSC-A as it facilitated appropriate gating to increased granularity of ciliated cells. Other cell types were gated using histograms, fluorescent spread shown. B) BT-IV-AF488 versus NGFR-PE shows a lack of co-staining consistent with population identities. C). BT-IV –AF488 versus MUC5AC-BV605 shows MUC5AC is ciliated and non-ciliated cells. D) MUC5AC-BV605 versus NGFR-PE shows a lack of co-staining consistent with population identities.

Supplementary Figure 2 Normalized gene count data generated through Partek Flow ‘quantify to annotation model’ normalized using rlog. (B) Distance matrices calculated from normalized count data. PCA of normalized count data.

Supplemental Table 1: Annotated DESeq2 results for differential gene expression by lineage. Supplemental Table 2: Gene Ontology analysis of differentially expressed gene.

Supplemental Table 3: Raw data MagPix Cytokine panel output with standard curves and analysis of geometric means. Supplemental Table 4: GISIAD isolate IDs and metadata for all sequences analyzed in this study.

Supplemental Table 5: Post-alignment quality control tables from RNAseq experiments.

## Author Contributions

Conceptualization, A.P., J.W.; Methodology, J.W., E.A., R.Z., A.J., A.D., H.L.;, Software, E.A., A.J., A.D.;, Validation, E.A., J.W.; Formal Analysis, J.W., E.A., A.J., A.D.;, Investigation A.P., J.W., E.A.;,.; Resources, A.P., K.F.., R.R…; Data Curation, E.,A.. A.J., A.D.; Writing – Original Draft Preparation, J.W., E.A., A.P.;, Writing – Review & Editing, J.W., E.A., A.P..; Visualization, J.W., E.A., A.P.,; Supervision, A.P. Project Administration, A.P..; Funding Acquisition, A.P.. All authors have read and agreed to the published version of the manuscript.

## Institutional Review Board Statement

IBV Clinical Isolate Collection: The human subjects’ protocol was approved by the Johns Hopkins School of Medicine Institutional Review Board (IRB90001667) and the National Institutes of Health Division of Microbiology and Infectious Diseases (protocol 15-0103).

Neutralizing Antibody Assays This study was approved by the JHU School of Medicine Institutional Review Board, IRB00288258. Serum samples for this study were obtained from healthcare workers (HCWs) recruited from the Johns Hopkins Centers for Influenza Research and Surveillance (JHCEIRS) during the annual Johns Hopkins Hospital (JHH) employee influenza vaccination campaign. Preand post-vaccination (∼ 28 day) human serum were collected from subjects, who provided written informed consent prior to participation.

## Informed Consent Statement

Informed consent was obtained from all subjects involved in the study. All human specimen were deidentified before use.

## Data Availability Statement

Raw FASTQ data for hNEC RNA sequencing can be found under bioproject: XXXXXX. All scripts used for subsequent analysis are available on GitHub under: https://github.com/Pekosz-Lab/IBV_transcriptomics_2023.

## Acknowledgments

We thank Tricia Nilles and Worod Allak from <u>the Becton Dickinson Immune Function Laboratory</u> at the Johns Hopkins Bloomberg School of Public Health, for flow cytometry training/support/technical assistance. The facility was supported in part by CFAR: 5P30AI094189-04 (Chaisson). We gratefully acknowledge all data contributors, i.e., the Authors and their Originating laboratories responsible for obtaining the specimens, and their Submitting laboratories for generating the genetic sequence and metadata and sharing via the GISAID Initiative, on which this research is based.

